# The evolution of structural variation across 500 million years of vertebrate evolution

**DOI:** 10.64898/2026.06.26.733778

**Authors:** Runyang Nicolas Lou, Daven Lim, Minoli Daigavane, Landen Gozashti, Gregory L Owens, Nilah M Ioannidis, The Vertebrate Genomes Project Consortium Phase I, Peter H Sudmant

## Abstract

Structural variants (SVs) contribute substantially to genetic variation and play vital roles in adaptation and disease^1,2^. Nonetheless, SVs are poorly captured by short reads and thus remain understudied, especially in non-model organisms^3,4^. Here, using haplotype-resolved genome assemblies from >600 vertebrate species, we comprehensively survey the landscape of SVs across >500 million years of evolution. We identify 35.3 million SVs and 3.12 billion single nucleotide variants (SNVs) segregating between two representative haplotypes across species, with SVs impacting ∼12-fold more base pairs. SV and SNV heterozygosity are correlated across species, with endangered and threatened species exhibiting reduced genetic diversity. However, the contribution of SVs relative to SNVs fundamentally differs across major vertebrate clades: given the same number of SNVs, fishes, amphibians, and reptiles have 4.3-to-9.1 times the number of SVs than birds, and 1.7-to-3.6 times more than mammals. This reduction in the relative contribution of SVs in mammals and birds is linked to fewer non-repeat-associated SVs as well as lower transposable element (TE) abundance and diversity. We identify features underlying genomic instability across vertebrates, finding that SVs frequently occur in repetitive and SNV-rich regions and are mediated by both homology and non-canonical DNA structures. Notably, G-quadruplex structures are enriched 11.5-fold around SV breakpoints in birds, while Z-DNA structures are enriched 2.2-fold in cartilaginous fishes. TEs uniquely contribute to SVs both directly through transposition and indirectly by mediating ectopic recombination, with the proportion of TE-mediated SVs influenced by both genomic TE density and diversity. We identify >10,000 instances of recent TE turnover including extinction of LINE-2 in therian mammals and slowing of CR1 activity in passerine birds. Finally, we show that SVs have an outsized role in functional genetic variation and are >70 times more likely to strongly impact protein-coding sequences than SNVs. While SVs are on average deleterious, we identify extensive recurrent structural variation across multiple taxa in genes involved in sensory, immune, and metabolic systems. Together, this study highlights extraordinary variation in the abundance, composition, mechanism, and functional impact of SVs across vertebrates.

## Main

Genetic diversity is the fundamental substrate of all phenotypic diversity on this planet^5^. Understanding the molecular and evolutionary forces that drive and maintain different types of genetic diversity across taxa is thus a central goal of evolutionary biology^6,7^. Over the past few decades, advances in high-throughput sequencing approaches have transformed our understanding of single nucleotide variants (SNVs), demonstrating that both the mutation rate and spectrum vary substantially across diverse taxa, often shaped by various life history traits as well as environmental factors^8–13^. Structural variants (SVs), including insertions, deletions, duplications, inversions, and complex rearrangements, are another important contributor to genetic diversity^14^. SVs exhibit drastic variation in scale spanning from 50 bp to chromosome-scale rearrangements^14^. Extensive studies in humans and other model systems have shown that SVs tend to form in repetitive regions of the genome and that transposable element (TE) activity is a key driver of SV formation^15–17^. Given the heterogeneity of SVs, they can potentially exhibit vast variation in their contributions to genetic diversity, length distributions, and mechanisms of formation across different species^17,18^. Importantly, SVs can introduce large-scale, large-effect changes to the genome instantaneously with important functional implications^1,19,20^ and are increasingly appreciated for their roles in adaptation^21–24^, speciation^25,26^, and disease^27,28^ across diverse organisms.

However, SVs have proven exceptionally challenging to assay with short reads due to their length and often complex genomic substrates^3^. Thus, outside of a few model systems^17,18,29,30^, our understanding of SVs and their contributions to diversity and evolutionary processes is extremely sparse and often unsystematic. Recent advances in long-read sequencing technologies^31^ and genome assembly methods^32,33^ have enabled accurate reconstructions of both haplotypes of diploid individuals, including structurally complex regions. By aligning these haplotypes against each other, both SNVs and SVs that segregate within an individual can be confidently identified^3,14^, enabling the complete characterization of the full spectrum of genetic diversity (**Fig. 1A**). Here, we take advantage of high-quality haplotype-resolved assemblies generated as part of the Vertebrate Genomes Project (VGP)^34,35^ Phase I and others^36,37^ to comprehensively survey the diversity of SVs across 629 vertebrate species spanning over 500 million years of evolution^38^. Together, our results reveal fundamental differences in the abundance, composition, mechanistic basis, and impact of different forms of genetic diversity across vertebrate clades.

**Figure 1.**
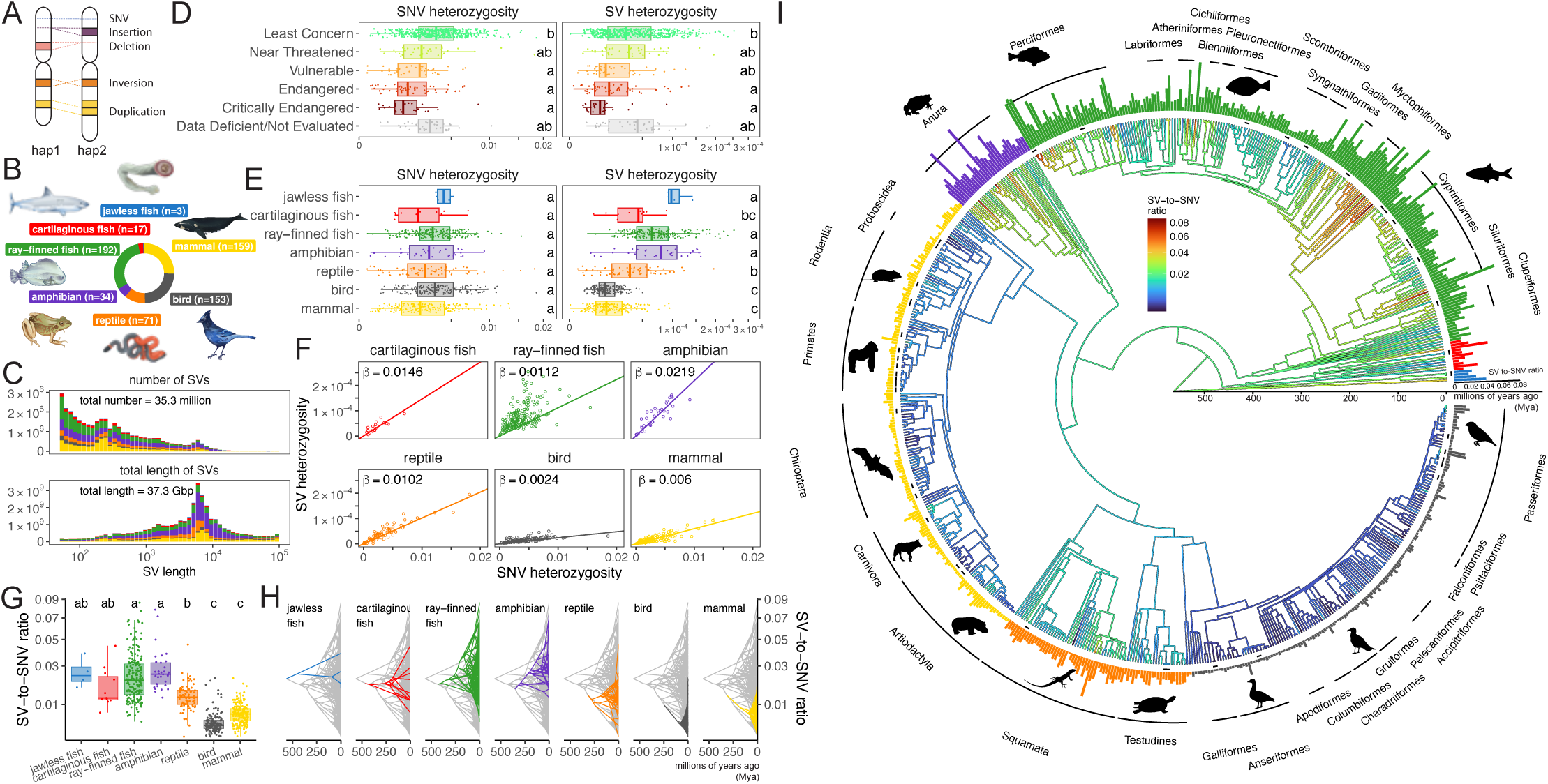
Distribution of SNV and SV diversity across the vertebrate tree of life. **A)** A conceptual schematic of our method for identifying SNVs and SVs from haplotype-resolved genome assemblies. **B)** The number of unique species included in our dataset, grouped by major vertebrate clade. **C)** The combined length distribution of all SVs that we identified, by number of variants (top) and number of base pairs impacted (bottom). The color scheme is the same as in **B)**. **D-E)** The distribution of SNV and SV heterozygosity grouped by IUCN conservation category and by clade. Different letters on the right indicate statistically significant differences between categories (Bonferroni-adjusted p-value threshold of 0.05, phylogenetic generalized least squares followed by Tukey’s test). **F)** The correlation between SV and SNV heterozygosity grouped by clade. The regression coefficient is indicated in each plot (phylogenetic generalized least squares with fixed intercept). **G)** The distribution of SV-to-SNV ratio grouped by clade. **H)** The ancestral state reconstruction of the SV-to-SNV ratio (shown on the y axis) along the vertebrate phylogeny. Each major clade, including the branch leading to it, is highlighted in its own panels while the rest of the tree is shown in grey. **J)** The ancestral state reconstruction of SV-to-SNV ratio presented in a circular tree, with colors corresponding to the reconstructed trait value. Bars at the tip represent the SV-to-SNV ratio in each assembly, colored by clade. Orders with more than four assemblies are labeled. Multiple assemblies of the same species are indicated with short lines next to their tips.

### A catalog of structural and single nucleotide diversity across 674 haplotype-resolved vertebrate genomes

We collated haplotype-resolved assemblies of vertebrates from NCBI and performed stringent quality filtering for highly contiguous and complete genomes, resulting in a total of 674 genomes from 629 unique vertebrate species (**Fig. 1B**, **Extended Data Fig. 1A**, **Table S1**). All assemblies are derived from PacBio long-read data, with some having supplemental Illumina short-read, ONT, and Hi-C data. These genomes are highly contiguous across both haplotypes (contig N50 of both haplotypes >100 kbp in all assemblies, >1Mbp in 471 assemblies) (**Extended Data Fig. 1B-C**). Collectively, these assemblies capture a vast amount of vertebrate diversity, with all major extant lineages well-represented except for the lobe-finned fishes (i.e. lungfish and coelacanth) (**Fig. 1B**). The majority (n=476) of these genomes were generated by the VGP^35^, with additional contributions from the California Conservation Genomics Project (CCGP)^36^, the Canadian Biogenome Project (CBP), and other consortia and individual institutions (**Table S1**). To characterize genetic diversity, we aligned both haplotypes to each other and jointly called SNVs, small indels (1-49 bp), and SVs (50 bp-100 kbp) using the primary haplotype or more contiguous haplotype as an arbitrary “reference”. In total, we cataloged 35.3 million SVs spanning 37.3 Gbp (**Fig. 1C**) along with 3.12 billion SNVs and 618 million small indels (see Methods for details). Together, these data represent the largest ever catalog of SV and SNV genetic diversity among vertebrate species to our knowledge. The vast majority of SVs are insertions and deletions, with inversions and duplications representing <0.2% of all SVs identified (**Table S2**). This likely reflects both the lower frequency of these variant types as well as technical challenges in their identification^3^. We thus focused our analyses on insertions and deletions.

### A fundamental shift in the contribution of structural variation to genetic diversity during vertebrate evolution

To quantify the contribution of different forms of variation to genetic diversity across clades, we calculated heterozygosity (i.e. the density of genetic variants along the genome) for both SNVs and SVs (**Fig. 1D-E, Extended Data Fig. 2A-C, Table S2**). Both SNVs and SVs heterozygosity were significantly correlated with IUCN conservation status as both are shaped by demography, highlighting that a single haplotype-resolved reference genome can be a valuable resource for assessing species biodiversity (**Fig. 1D, Extended Data Fig. 2A,** see Supplementary Notes and Formenti et al.^35^). SNV heterozygosity (HSNV) varies by over two orders of magnitude across vertebrate species, with more variation observed within clades than between clades (**Fig. 1E, Extended Data Fig. 2C)**. SV heterozygosity (HSV) similarly spans two orders of magnitude across species. Although HSV is ∼100 times lower than HSNV on average, the total number of base pairs impacted by heterozygous SVs is ∼10-fold more than heterozygous SNVs (**Extended Data Fig. 3A**). In stark contrast to HSNV, however, HSV exhibits extensive between-clade variation. Jawless fish, ray-finned fish, and amphibians have significantly higher HSV than birds and mammals, with reptiles intermediate (Bonferroni-adjusted p-values<0.0009, phylogenetic generalized least squares followed by Tukey’s test, see Methods for details, **Fig. 1E**). In fact, the highest HSV found among 153 bird species (Acorn woodpecker, *Melanerpes formicivorus*, HSV=5.02×10^-5^) is lower than the median values in ray-finned fish and amphibians (6.38×10^-5^ and 7.77×10^-5^, respectively).

To explore the extensive variation in SV diversity between clades, we contrasted SNV and SV heterozygosity within each clade. As expected, HSNV and HSV are strongly positively correlated within clades due to the influence of demography and effective population size on these statistics (p-values<0.005, phylogenetic generalized least squares) (**Fig. 1F**). However, the regression coefficients (β) differ vastly among major vertebrate clades. Thus, given the same number of SNVs, fishes, amphibians, and reptiles tend to have 4.3-to-9.1 times the number of SVs than birds, and 1.7-to-3.6 times more than mammals.

We calculated the ratio of HSV to HSNV in each assembly to quantify the relative contribution of SVs to genetic diversity in individual species (the SV-to-SNV ratio, **Fig. 1G, Extended Data Fig. 2D**). This statistic exhibited strong phylogenetic signal (Pagel’s λ=0.983, p-value=2.8×10^-126^) across vertebrates and an ancestral state reconstruction demonstrated that the SV-to-SNV ratio has experienced several major declines throughout vertebrate evolution (**Fig. 1H-I**). While the most recent common ancestor (MRCA) of all vertebrates exhibited 0.022 SV for every SNV, this declined to 0.014 in the MRCA of all extant amniotes (reptiles, birds, and mammals) . Subsequently, the SV-to-SNV ratio further declined independently in the MRCAs of mammals (0.0089), crocodilians (0.0055), and birds (0.0046) (**Fig. 1H-I, Extended Data Fig. 4**). The resulting landscape of genetic diversity is fundamentally different across taxa: diversity in fish and amphibians is more extensively shaped by structural variation in comparison to mammals, birds, and reptiles.

Though the composition of genetic diversity is relatively consistent within each vertebrate clade, outliers were observed in addition to these broad-scale patterns. Some notable examples of increases in the relative contribution of SVs include ray-finned fish in orders Gadiformes, Myctophiformes, and Perciformes, some New World passerine birds of the families Icteridae and Passerellidae, and the island scrub jay (*Aphelocoma insularis*) (**Fig. 1I, Extended Data Fig. 4**). Clades that show decreases in their SV-to-SNV ratio include ray-finned fish of the order Scombriformes and Syngnathiformes, and bats of the family Phyllostomidae.

Sometimes sharp increases in the contribution of SVs are followed by declines in a subclade. For instance, the MRCA of cats (of family Felidae) and cetaceans (of infraorder Cetacea) both experienced elevated SV-to-SNV ratios, resulting in species exhibiting some of the highest ratios observed in mammals. These include the clouded leopard (*Neofelis nebulosa*, 0.022), the cougar (*Puma concolor*, 0.015), the Northern bottlenose whale (*Hyperoodon ampullatus*, 0.018), the Sowerby’s beaked whale (*Mesoplodon bidens*, 0.015), and some other cetaceans (**Table S2**). Such surges, however, were subsequently dampened in the genus *Panthera* and the family Delphinidae. Together, these results demonstrate that the composition of genetic diversity is fundamentally different among vertebrate clades and species, with both broad-scale changes in deep evolutionary times as well as frequent turnovers in shorter time scales.

### Taxon-specific drivers of SV diversity

Structural variants are often made up of repetitive sequences such as transposable elements (TEs), satellite repeats, and simple repeats. Vertebrate genomes vary dramatically in both the relative abundance and genomic distribution of these repeat types^39,40^. We quantified the composition of SVs across taxa by classifying them into five categories: TE, simple repeat, satellite repeat, other repeat (e.g. RNA arrays, low complexity repeat), and non-repeat (**Fig. 2A left panel,** see Methods and Supplementary Notes for details). If an SV cannot be unambiguously assigned to a single repeat type, we classify it as a non-repeat SV. TE, simple repeat, and non-repeat sequences are the most common SV types, together representing 97.5-99.4% of SVs on average in all major vertebrate clades. However, these repeat classes vary extensively across clades: TEs constitute the majority of SVs in mammals, reptiles, and amphibians, making up 51.2-60.1% of SVs on average in each of these clades (**Fig. 2B left panel**). In contrast, in fishes and birds, non-repeat SVs are the most common, making up 43.7-52.3% of SVs. Simple repeats are consistently the third leading contributor to SVs, underlying 4.2-29.9% of SVs in each clade on average and up to 60% in individual species. In several primate and bird species, satellite repeats constitute another significant component of SVs (up to 38%), although this partially reflects difficulties in assembling and annotating satellites.

**Figure 2.**
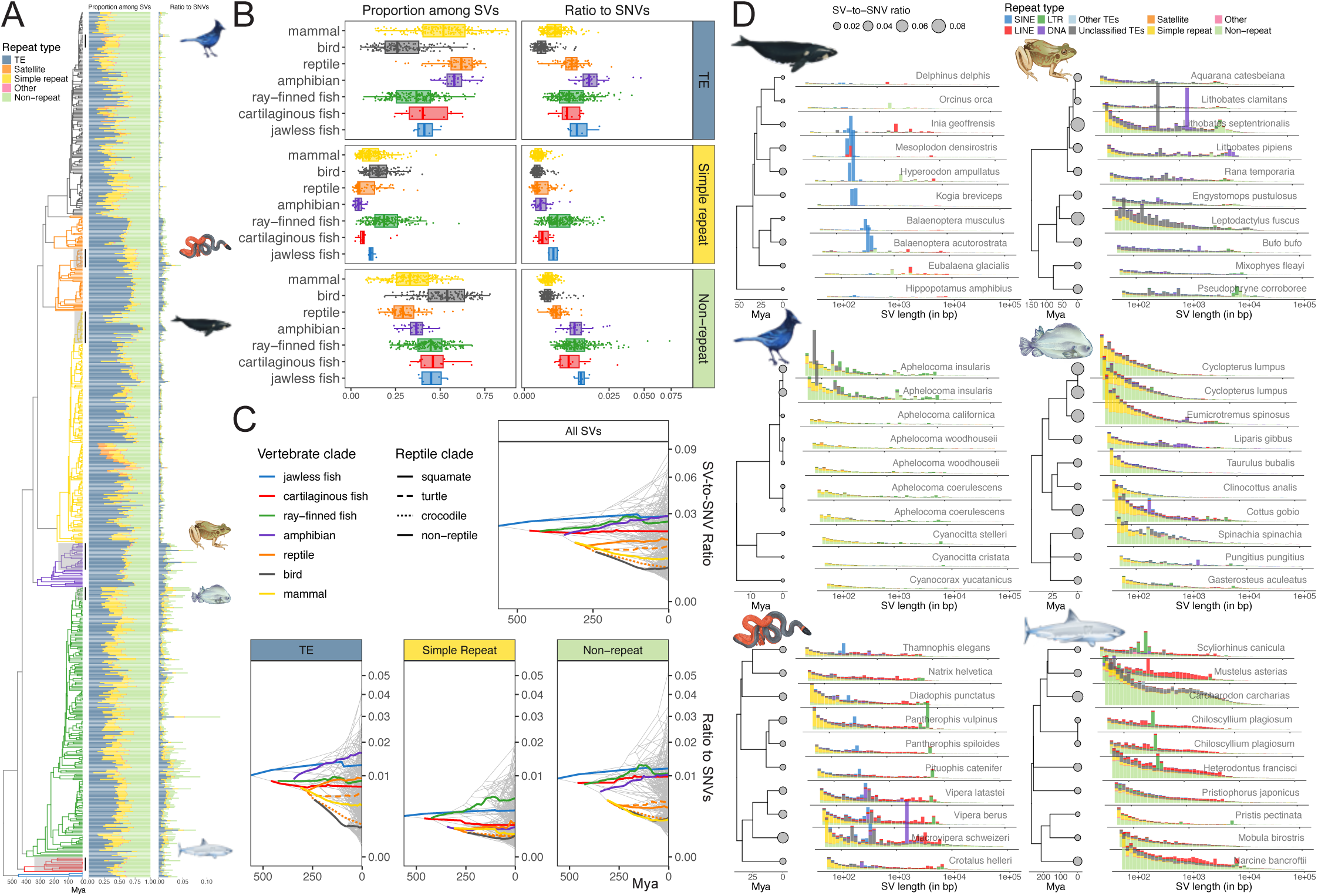
Taxon-specific drivers of SV diversity. A-B) The distribution of SV types by their proportion among all SVs (left column) and by their contribution to the SV-to-SNV ratio (right column) along the vertebrate phylogeny in **A)** and grouped by major clade in **B)**. Highlighted subclades in **A)** contain examples of rapid SV-to-SNV ratio turnovers and are further examined in **D)**. **C)** The ancestral state reconstruction of SV-to-SNV ratio shown on top and broken down by SV types at the bottom. The average reconstructed trait values for each major clade (including the branch leading to it) over time are shown in colored lines. The three reptile clades are distinguished by different line types. **D)** Highlighted examples of SV composition and length distribution in six subclades. In each example, the size of the circles at the tips of the phylogeny indicates the SV-to-SNV ratio in each assembly. To their right, the histograms show the length distribution of SVs normalized by SNVs, colored by SV type. Note that the x-axis in each example is slightly offset to reduce overlap between plots.

We next asked if these differences in the composition of SVs explained the reduced SV-to-SNV ratios in amniotes by decomposing the ratio into different SV types (**Fig. 2A-B right panel**) and performing ancestral state reconstruction for each SV type separately (**Fig. 2C, Extended Data Fig. 5A-C**). In fishes, non-repeat and TE SVs both contribute substantially to total genetic diversity, suggesting that the ancestral state of high SV-to-SNV ratios in vertebrates was likely driven by high TE activities and an abundance of non-repeat SVs. The relative contribution of TEs subsequently underwent additional increases in amphibians, remained relatively steady in squamates and turtles, and independently decreased in mammals, crocodilians, and birds. The contribution of non-repeat SVs to genetic diversity decreased in the MRCA of amniotes and further in mammals, crocodilians, and birds. Contributions of simple repeats are relatively low in ancestral vertebrates but have independently increased in ray-finned fishes and snakes. Thus, the reduced SV-to-SNV ratio in the amniote ancestor is mainly explained by fewer non-repeat SVs while further decreases in mammals, crocodilians, and birds are driven by both additional reductions in TEs and non-repeat SVs.

Within clades, SV-to-SNV ratio outliers were also driven by different types of SVs. For example, the expansions and contractions of SV-to-SNV ratio in cetaceans and cats are mainly due to the activation and suppression of SINE/tRNA elements in these subclades (**Fig. 2D, Extended Data Fig. 3B-D**). This elevated SINE/tRNA activity results in a large proportion of ∼300-bp SVs in the distribution of SV sizes. Similar peaks driven by the transposition of DNA/TcMar-Tc1 elements are observed in the Milos viper (*Macrovipera schweizeri*) and the mink frog (*Lithobates septentrionalis*), two species that exhibit heightened SV-to-SNV ratios compared to their close relatives. In contrast, non-repeat and simple-repeat SVs underlie increased SV-to-SNV ratios in other species, including the island scrub jay (*Aphelocoma insularis*), the foxsnake (*Pantherophis vulpinus*), the rufous frog (*Leptodactylus fuscus*), lump fishes (of family Cyclopteridae), and the great white shark (*Carcharodon carcharias*). Together, we find that TEs dominate the SV-to-SNV ratio evolution within mammals, explaining >75% of the variance among mammalian species (**Extended Data Fig. 5D**). Non-repeat SVs explain ∼70% of the variance in cartilaginous fishes, while all three SV types contribute more evenly in other clades. These results demonstrate highly punctuated and unsynchronized contributions of different forms of structural variation to diversity both within and between major vertebrate clades.

### Frequent turnover of TE composition in SVs and genomes

TEs are among the top two contributors to SVs in all major vertebrate clades. However, TEs exhibit extensive diversity in their origins, mechanism of replication, and sequence features. Broadly, they can be grouped into two classes: Class I TEs (i.e. retrotransposons) mobilize through an RNA intermediate; Class II TEs (i.e. DNA transposons) do not. Each class of TEs is further divided into subclasses and families based on their characteristics (**Fig. 3A**). While TE diversity has been extensively characterized in genome assemblies of diverse species, insertions that are still segregating within a species (i.e. TE SVs) provide a unique window into recently active TEs, which are particularly relevant to contemporary evolutionary processes.

**Figure 3.**
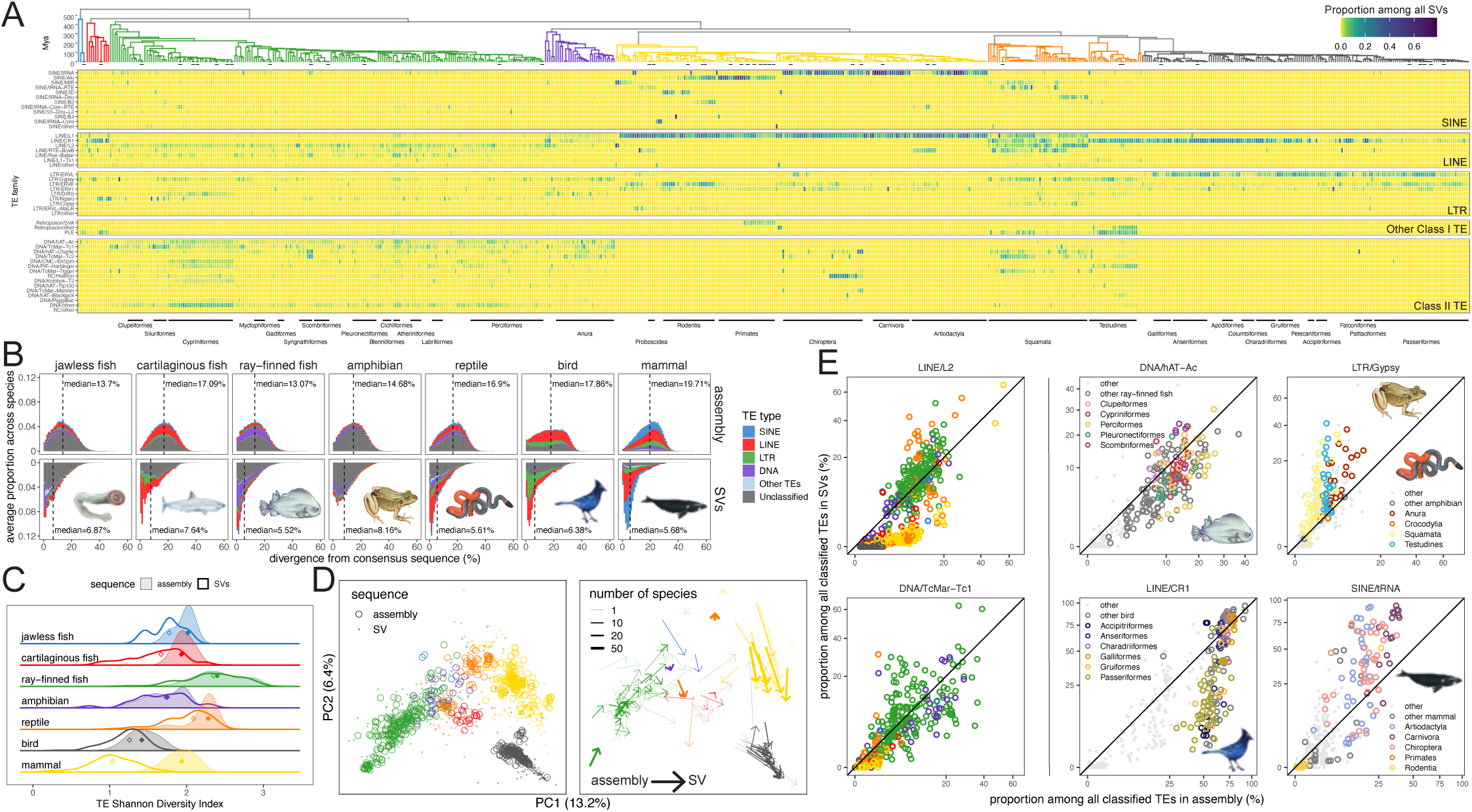
TE composition in SVs and genome assemblies. **A)** Heatmap of TE composition in SVs along the vertebrate phylogeny. Each column represents an assembly, and each row represents a TE family. The color of each cell corresponds to the proportion of SVs assigned to a given TE family out of all SVs in each assembly. Major vertebrate orders are labeled at the bottom of the heatmap. **B)** The distribution of divergence from the consensus sequence for TEs in genome assemblies (top) and SVs (bottom), grouped by clade and colored by major TE orders. The median divergence value is annotated in each plot. **C)** The distribution of Shannon Diversity of TEs in genome assemblies (filled density curves) and in SVs (empty curves), grouped by clade. Diamond-shaped markers indicate median values. **D)** A PCA of TE composition (i.e. proportions of TE families among all classified TEs) in genome assemblies and SVs, colored by clade. In the left panel, assemblies are shown in circles and SVs in dots. In the right panel, an arrow is drawn to represent the average shift in TE composition in SVs versus assemblies in each vertebrate order. The width of each arrow corresponds to the number of assemblies in the order. **E)** The abundance of six highlighted TE families in SVs (y-axis) and in genome assemblies (x-axis), as quantified by their proportion among all classified TEs. Assemblies are colored by major clades in the first two examples, and by major orders in ray-finned fishes, birds, amphibians and reptiles, and mammals in the last four examples, respectively. A one-to-one line is drawn to indicate the expected distribution if TE composition is identical between SVs and assemblies.

We classified 10.5 million TE SVs into 130 TE families, constructing a vertebrate-wide catalogue of recent mobile element activity (**Supplementary Data 1)**. These segregating TE SVs are significantly younger than non-segregating TEs in assemblies, exhibiting lower divergence from their consensus sequences (**Fig. 3B**). The distribution and diversity of these recently active TEs show extensive variation across vertebrate species (**Fig. 3A, C**). Notably, ray-finned fishes have the highest Shannon Diversity in their TE SV composition, potentially driven by extensive horizontal transfer of TEs found in this clade^41^. Jawless fishes, cartilaginous fishes, amphibians, and reptiles similarly exhibit a broad spectrum of different TE types contributing to SVs. Diversity of TE SVs in birds is the lowest, consisting mostly of LINE/CR1 elements and several LTR families while largely devoid of Class II TEs. Mammals similarly lack Class II TEs (with the exception of bats) while exhibiting higher SINE and LINE diversity and activity.

In addition to segregating SVs resulting from recent TE insertions, genomes are riddled with ancient TE insertions providing a record of historical TE activity. Contrasting the composition of TEs in the assembly with more recently active segregating TEs allows us to investigate TE evolution and turnover through time. TEs in SVs are a subset of TEs in the genome and therefore are less diverse (**Fig. 3C**). Mammals in particular have a much smaller set of segregating TEs compared to TEs in their genomes, consistent with a more diverse TE repertoire in the mammalian ancestor^42^. We performed a PCA of TE composition with both genomic and segregating TEs to characterize changes in TE composition throughout evolution (**Fig. 3D**). The TE composition of the genome exhibits strong phylogenetic signal in PC space. Segregating TEs largely mirror the distribution of genomic TEs. However, many species exhibit marked shifts indicating recent changes in TE activity. Several of these changes were concordant within clade (e.g. in birds and mammals), likely reflecting shared TE turnover in their recent common ancestor. To identify which TE elements are driving these patterns we compared the relative proportions of different TE families in genome assemblies versus SVs. We identify >10,000 instances of significant shifts in TE activity across species (Fisher’s exact test followed by Bonferroni correction, p-value threshold of 0.05, **Supplementary Data 1, Fig. 3E, Fig. S1-6**). For instance, LINE/L2 elements are present in substantial proportions in mammalian genomes. On average, they represent 5.1% of all classified TEs in eutherian mammals, 18.9% in marsupials, and 52.6% in monotremes. However, they are mostly absent from the SVs of therian mammals (0.2% in eutherians, 2.4% in marsupials, and 52% in monotremes), consistent with previous finding that LINE/L2 were active in the mammalian ancestor but subsequently went extinct in therians^43^. In birds, LINE1/CR1 elements are the most abundant TE family in all but three genomes (62.8% on average), yet their contribution to SVs is substantially lower in many subclades including most passerines (53.7% in genomes, 18.4% in SVs), consistent with lineage-specific suppression. In contrast, in mammals of the superorder Laurasiatheria (including Artiodactyla, Canivora, Chiroptera, and others), the proportions of SINE/tRNA elements in SVs are highly variable and often far outweigh their share of the genome (18.6% in genomes, 40.7% in SVs on average), consistent with shared expansions of SINE/tRNA elements in the MRCA of these clades followed by frequent turnovers in subclades (**Fig. 2D, Extended Data Fig. 3D**). Together, these results show that turnover in TE activity is highly dynamic in vertebrate species which in turn can dramatically shape the SV mutational landscape and that SVs are a valuable resource for exploring TE activity across species over time^44,45^

### The genomic substrate and mechanisms underlying SV formation

The formation and maintenance of SVs in the genome is strongly shaped by both local genomic features and sequence content. To explore the relative contributions of different genomic features to SVs across taxa, we took a predictive modeling approach. We trained a random forest model in each species to predict the presence of SVs using nearby genomic annotations (excluding SV sequences themselves, **Fig. 4A**, see Methods for details). This type of model has previously been shown to exhibit strong predictive power in humans (AUROC=0.904)^46^. Despite having substantially smaller sample size and fewer annotated genomic features than its application in humans, the model performed well across diverse vertebrate species, with an AUROC of 0.81 on average (ranging between 0.697 and 0.975) (**Extended Data Fig. 6A**). The top predictive features identified are similar across clades (**Fig. 4A, Extended Data Fig. 6B**). The local density of SNVs is consistently the most important feature, showing strong positive correlation with SV presence, reflecting shared mutational mechanisms and constraints between SNVs and SVs (**Extended Data Fig. 6C**). Distance to nearby repetitive sequences similarly contributes substantially to the predictive power across clades. This suggests shared underpinnings of genomic instability in vertebrates, where SVs are more commonly found in repetitive, SNV-rich regions. However, we also identify differences across clades revealing variation in the mechanistic basis of SV formation (**Fig. 4A**). For example, distance to simple repeats is the second most predictive feature in ray-finned fish, consistent with our observation of increased simple repeat SVs in this clade (**Fig. 2B,C**). Similarly, distance to SINE elements is strongly predictive of SVs only in mammals, reflecting their elevated SINE element activity (**Fig. 3A**). Moreover, high GC content is strongly predictive of SVs in cartilaginous fishes and archelosaurs (turtles, crocodilians, and birds) (**Extended Data Fig. 6B-C**).

**Figure 4.**
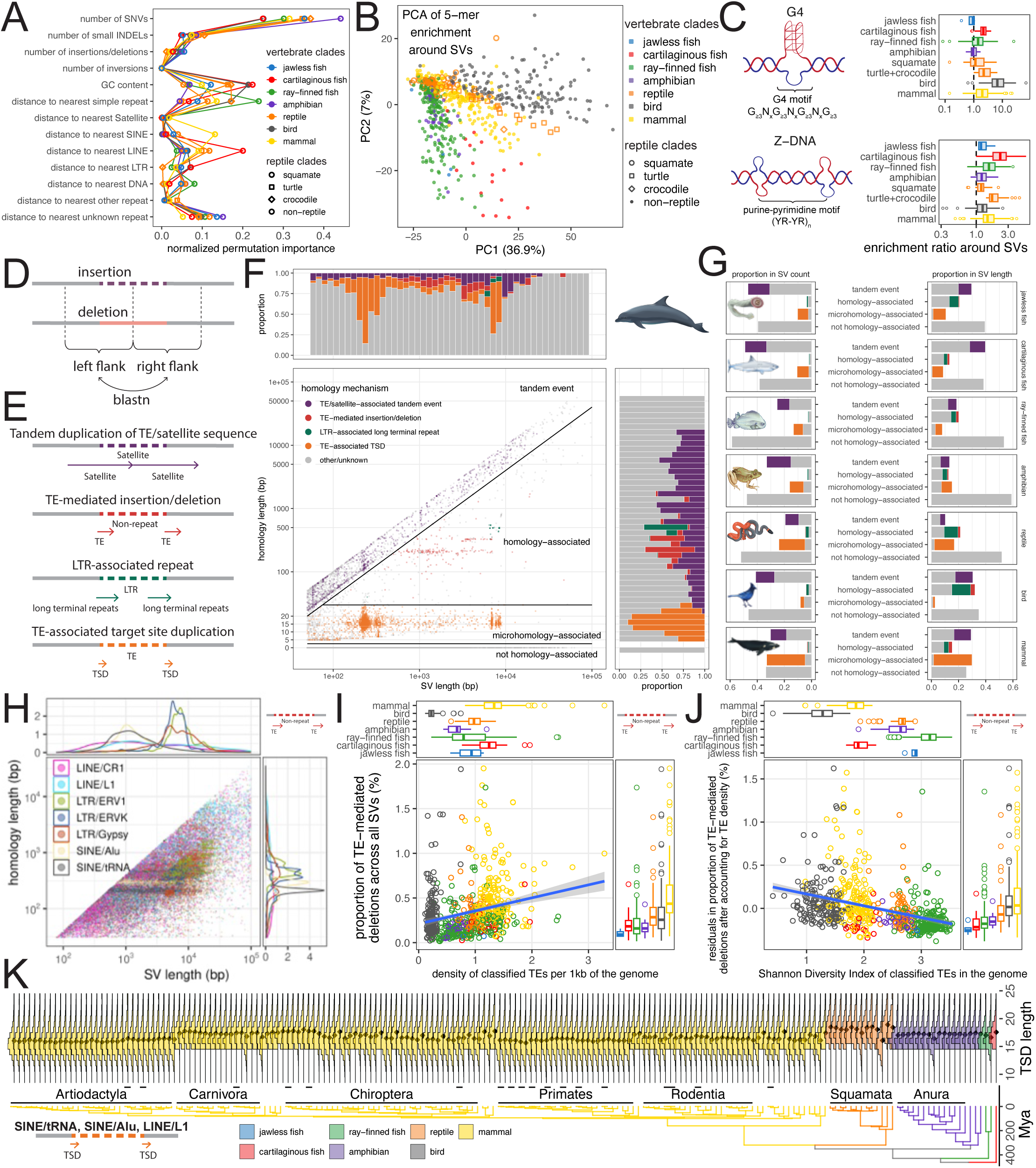
The genomic substrate and mechanisms underlying SV formation. **A)** The normalized permutation importance of genomic features in predicting SV presence through a random forest model. The average importance score is shown for each clade, differentiated by colors and point shapes. **B)** PCA of 5-mer enrichment ratios in 400-bp windows around SV breakpoints. **C)** The enrichment ratio of G4- and Z-DNA-associated motifs in 400-bp windows around SV breakpoints, grouped by clade. **D)** A conceptual schematic of our method for identifying homologous sequences spanning SV breakpoints. Note that insertion sequences were reinserted into the reference before the homology search. **E)** Examples of homology-forming mechanisms around SV breakpoints. **F)** An example showing the relationship between the length of SVs and the length of homology sequences at SV breakpoints in common bottlenose dolphin (*Tursiops truncatus*). The plot is divided into four zones, illustrating our categorization of different homology patterns. Specific homology-forming mechanisms can sometimes be identified based on the repeat annotation of SVs and homology sequences, as shown by colored points. Marginal plots show the SV and homology length distributions colored by these mechanisms. **G)** The proportion of SVs with each homology category in each clade, quantified by the number of SVs (left) and the number of base pairs impacted (right). Colors correspond to cases where homology-forming mechanisms can be identified as in **F)**. **H)** The distribution of SV length and homology length in insertions/deletions mediated by select TE families, indicated by different colors, across all vertebrate species. **I)** The relationship between the proportion of TE-mediated insertions/deletions (y-axis) and TE abundance (quantified as the density of classified TEs in the genome assembly, x-axis), colored by clade. A linear regression line is shown (p-value<2×10^-16^). **J)** The relationship between the residuals in TE-mediated insertions/deletions and the Shannon Diversity of TEs in the assembly, after taking the effect of TE abundance into account. A linear regression line is shown (p-value<2×10^-16^). **K)** The evolution of target site duplication (TSD) length associated with LINE1 endonuclease along the vertebrate phylogeny.

To further explore clade-specific GC bias surrounding SVs, we contrasted genome-wide sequence content to that around SV breakpoints using 5-mers. The enrichment of 5-mers around SVs exhibited phylogenetic signal in PCA space (**Fig. 4B, Extended Data Fig. 6D**). Examination of PC loadings revealed that an enrichment of GC-rich 5mers, such as TCCCC and GGGGC, separates cartilaginous fishes and archelosaurs from other clades along PC1 (**Fig. 4B, Extended Data Fig. 6E-F**). We hypothesized that these GC-rich k-mers might be indicative of sequences capable of forming G-quadruplexes (G4), non-canonical DNA structures that are unstable and prone to structural mutations^47^. In silico annotation of G4-associated motifs showed strong enrichment at SV breakpoints in cartilaginous fishes, turtles, crocodilians, and particularly in birds (11.5-fold enrichment on average), thus suggesting that they likely play an important role in SV formation in these clades (**Fig. 4C, Extended Data Fig. 6G-H**). We similarly identified enrichments of 5mers with altering purine-pyrimidine patterns (e.g. GTGTA, TACAC) separating cartilaginous fishes from other clades along PC2 (**Fig. 4B, Extended Data Fig. 6E-F**). These 5-mers are associated with Z-DNA, another unstable non-canonical DNA structure^47^. We find that Z-DNA motifs are modestly enriched at SV breakpoints in all vertebrate clades but especially in cartilaginous fishes (2.2-fold enrichment on average compared to 1.4-fold in other clades, **Fig. 4C, Extended Data Fig. 6G-H**). Together, these findings highlight non-canonical DNA structures as a major contributor to clade-specific differences in SV mutational mechanisms.

Local sequence homology in the genome can often lead to replication errors and ectopic recombination driving SV formation. Some TEs also introduce homologous sequences to their insertion sites, either through target site duplication (TSD) or long terminal repeats associated with LTR elements^48^. We thus searched for stretches of homology across SV breakpoints (**Fig. 4D**) following a similar workflow as described in Schloissnig et al.^29^. SVs were then classified into four types based on their homology patterns: non-homology-associated, microhomology-associated, homology-associated, and tandem events (**Fig. 4E-F**). We further annotated the homologous sequences based on their repeat content. We find that tandem events and non-homology-associated SVs are the two most abundant categories in most clades (**Fig. 4G**). Tandem events often involve TEs and satellites (40.7% of all tandem events), though a large fraction of non-repeat SVs also likely arose through this mechanism (40.3% of all non-repeat SVs, **Extended Data Fig. 7A**). Tandem events tend to be associated with smaller SVs (621 bp on average vs. 1,223 bp in other categories). Simple repeats, TEs that do not create TSDs, and non-repeats all contribute to the SVs that have no sequence homology across their breakpoints (**Extended Data Fig. 7A**). In contrast to other clades, reptiles and mammals have large contributions from microhomology-associated events due to TSDs left by TE insertions (**Fig. 4G**). Homology-associated SVs are the least abundant category across all taxa (3.4% of all SVs), although they tend to result in larger SVs (5,153 bp on average vs. 909 bp in other categories, **Fig. 4G**). In fact, many large non-repeat SVs likely arose due to ectopic recombination mediated by homologous sequences (e.g. 27.9% of all non-repeat SVs >10 kbp, **Extended Data Fig. 7B**).

Examination of the homologous sequences flanking SVs revealed that many are TEs. These SVs are thus likely the result of ectopic recombination between two TE copies, highlighting the varied ways in which TEs can drive genome instability. Across all taxa we find that different types of TEs mediate diverse patterns of SV formation (**Fig. 4H, Extended Data Fig. 7C**). SINE-mediated SVs tend to be driven by full-length copies of SINE elements (e.g. narrow peak in homology length corresponding to the length of Alu and tRNA elements). These SVs tend to be smaller (median=1,140 bp, interquantile range=693-2,087 bp). Partial LINE elements and class II TEs mediate SVs that vary widely in length (median=1,896 bp, interquantile range=692-6,499 bp). More broadly, the total proportion of TE-mediated SVs is positively correlated with TE content in the genome (p-values<2×10^-16^, linear model, **Fig. 4I, Extended Data Fig. 7D**). However, fishes, amphibians, and reptiles consistently exhibit fewer TE-mediated SVs than predicted by their genomic TE content. One reason for this observation could be that these genomes exhibit increased TE diversity, i.e. fewer homologous TEs are available to serve as substrate for SV formation (**Fig. 3C**). Consistent with this hypothesis, we find that after accounting for TE content, TE-mediated SVs are negatively correlated with the Shannon Diversity of TEs (p-values<2×10^-16^, linear model, **Fig. 4J**).

Our analyses of flanking sequence homology of SVs also identified variation in the length of TSDs at TE insertions across the phylogeny. Variation in TSD length reflects differences in TE endonuclease catalytic activity. TE families that rely on the same endonuclease exhibit conserved TSD lengths (**Extended Data Fig. 7E**). We find that TSD length variation is highly concordant with the phylogenetic tree for TEs that rely on the L1 endonuclease, with several lineage-specific shifts in the L1 endonuclease TSD length, including notable decreases in the TSD length in artiodactyls and increases in squamates. This is consistent with a history of mostly vertical transfer (**Fig. 4K**)^49^. In contrast, we find that the TSD length distribution is noisier along the tree for TEs that use the LINE/RTE-BovB endonuclease, consistent with frequent horizontal transfer (**Extended Data Fig. 7F**)^49^. Thus, TE evolution and activity reshape the distribution of genetic diversity across taxa in several varied ways.

### Functional impacts of SVs in adaptation and load across taxa

Structural variants often have large phenotypic effects due to their potential to disrupt functional genomic sequences. While SVs have been increasingly demonstrated to play outsized roles in adaptation, they are expected to be deleterious on average^19,20,27,50^. To investigate the potential functional impact of SVs across diverse vertebrate species, we analyzed 415 of our genomes from 413 species with existing gene annotations. We find that both SVs and SNVs become increasingly depleted across genomic regions of increasing functional importance, illustrating the impact of purifying selection (**Fig. 5A**). Notably, selection against SVs is far stronger than against SNVs, resulting in ∼2-fold lower enrichment ratios at exonic regions (**Fig. 5A**). This suggests that de-novo SVs are overall more deleterious than SNVs and are more efficiently purged across taxa, as has been previously reported in humans^18,51^.

**Figure 5.**
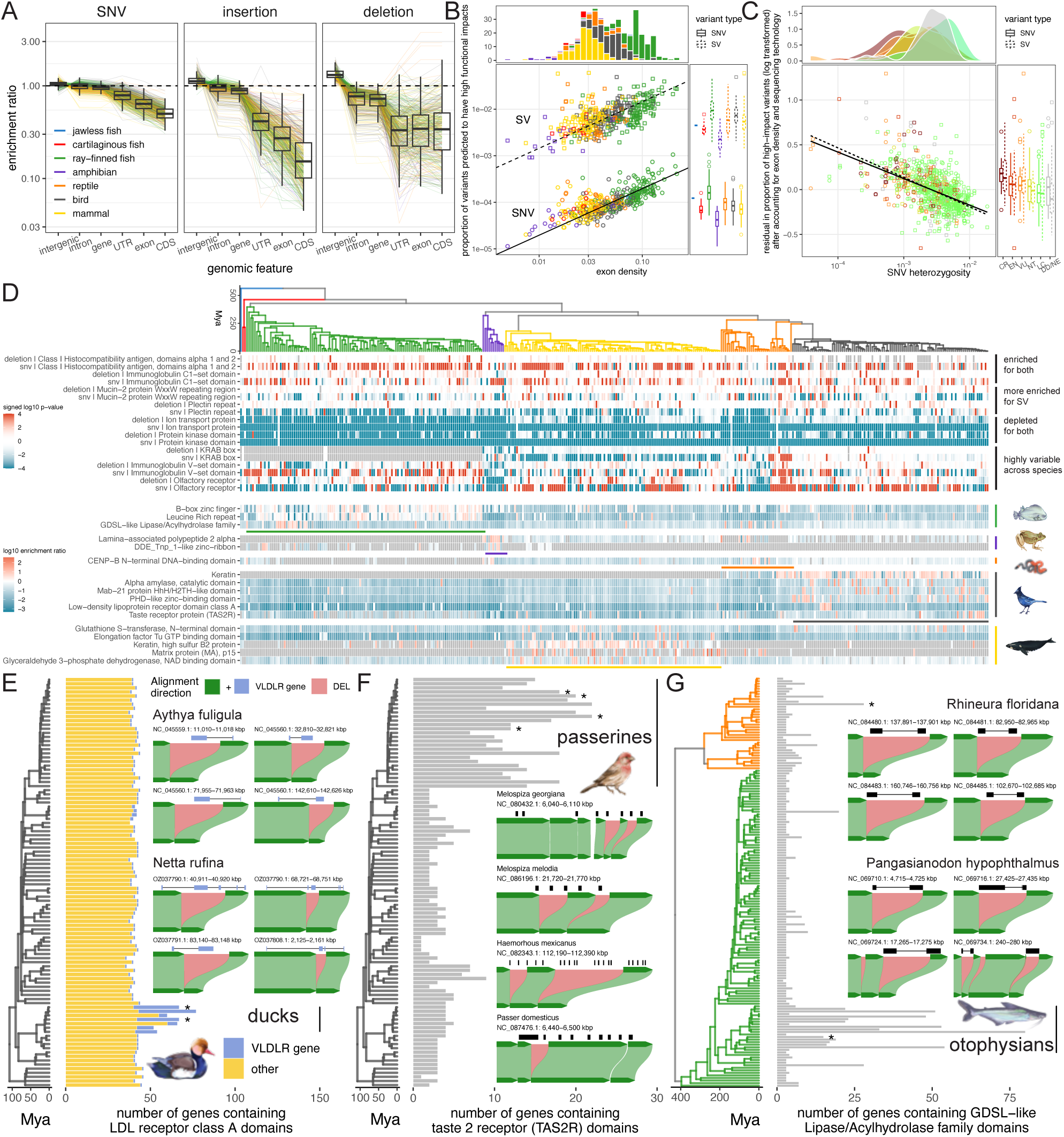
Functional impacts of SVs. **A)** Enrichment ratios of SNVs and SVs at genomic regions with increasing functional importance. Each assembly is shown as a line, colored by their clade, and the overall distribution is summarized in boxplots. **B)** The relationship between the proportions of SVs and SNVs predicted to have high functional impacts and exon density. SNVs are shown in circles and SVs are shown in squares. Alinear regression curve is fitted to SVs and SNVs separately, with sequencing technology (i.e. PacBio CLR vs. CCS) as a covariate in the model (p-values<2×10^-16^). Points are colored by clade. Marginal plots show the distribution of each variable broken down by clade. **C)** The relationship between residuals in the proportions of SVs and SNVs predicted to have high functional impacts and SNV heterozygosity, after taking the effect of exon density and sequencing technology into account. A linear regression curve is fitted to SVs and SNVs separately (p-values<2×10^-16^). Points are colored by IUCN conservation category. Marginal plots show the distribution of each variable broken down by IUCN category. **D)** The enrichment of select protein domains in their overlap with deletions and SNVs. The top heatmap shows examples of domains with relatively consistent patterns across all vertebrate species, broken down by those that are enriched with variants, depleted with variants, or show distinct patterns for deletions compared to SNVs. Colors of the heatmap indicate p-values testing for the significance of enrichment/depletion, capped at 10^-4^. The bottom heatmap shows examples of domains exhibiting clade-specific patterns in their intersection with deletions. Colors indicate enrichment ratio. In both heatmaps, enrichment is shown in red and depletion in blue. **E-G)** Three examples with protein domains showing clade-specific enrichment of deletions, all associated with clade-specific gene expansions. Bars at the tips of the tree indicate the number of gene copies that contain these protein domains in each species. Insets show examples of sequence alignment between two haplotypes in select species (indicated by stars in the tree) that harbor these deletions, leading to copy number differences between two haplotypes within a single individual. Exons are indicated by boxes while introns are represented by lines in gene tracks.

Given the strong purifying selection against de-novo SVs, we next compared the functional consequences of segregating SVs and SNVs on protein-coding genes using variant effect prediction^52^. We find that the proportion of variants with predicted high impact increases with exon density in the genome (p-values<2×10^-16^, linear model, **Fig. 5B**), because variants are more likely to intersect with protein-coding sequences in more compact genomes. However, given the same exon density, SVs are >70 times more likely to result in high impact mutations than SNVs (**Fig. 5B**). This finding highlights that despite being selected against and depleted at genic regions, SVs that persist are still far more likely to strongly alter genomic functions than SNVs by introducing large-scale changes. Accounting for the effect of exon density, we compared the proportion of variants with predicted high effects with SNV heterozygosity. We find that less diverse (and more endangered) species tend to harbor higher proportions of high-impact SVs and SNVs (p-values<2×10^-16^, linear model, **Fig. 5C**), consistent with higher genetic load in smaller populations and more efficient purging of deleterious variants in larger populations^20^.

Despite the potential deleterious consequences of SVs, these large-effect-size variants also have the potential to be adaptive. To identify putatively adaptive SVs, we searched for variants that have repeatedly altered the same genes across multiple species. Due to challenges associated with identifying orthologous genes across diverse taxa, we annotated protein domains across predicted gene sequences^53^ and tested for enrichment of SNVs and SVs. Since only the reference genome is annotated, the effect of insertions tends to be underestimated based on sequence overlap. We thus focused on deletions in this analysis. As shown previously (**Fig. 5A**), the vast majority of protein domains are depleted for genetic variation across all assemblies (**Extended Data Fig. 8A**). For example, many highly abundant domains such as those underlying ion transportation and protein kinases are highly conserved within each species (**Fig. 5D top panel, Extended Data Fig. 8A**). In contrast, domains that are known to exhibit structural diversity critical for immune functions, such as histocompatibility antigens and immunoglobulin domains, frequently harbor SVs and SNVs. Many repeat-containing domains, such as the mucin repeats and plectin repeats, are more enriched for SVs than SNVs, likely associated with copy number variation of these domains. Some domains, such as the olfactory receptor, are strongly depleted of genetic variation in some taxa and are strongly enriched for SVs and SNVs in others, potentially suggesting that these genes are under strong selective constraint in some taxa (e.g. amphibians and squamates) while undergoing extensive diversifying selection in others (e.g. mammals, turtles, and birds).

### Recurrent gene copy number variation across taxa linked to lineage-specific gene expansions

In addition to vertebrate-wide patterns, we also identified many instances of clade-specific enrichment of SVs in specific protein domains (**Fig. 5D bottom panel, Extended Data Fig. 8B-F**). Many of these are linked to clade-specific gene copy number expansions, with copy number variation potentially maintained by balancing selection within each species, highlighting the critical roles of gene duplications in adaptive evolution^54^. For example, ducks of the family Anatidae exhibit an excess of SVs in low-density lipoprotein receptor class A domains, encoded by both *LDLR* and *VLDLR* genes (**Fig. 5D**). While ducks have a comparable number of *LDLR* genes as other bird species, they experienced a drastic expansion of *VLDLR* genes (6-34 copies compared to 0-3 copies in other bird species, **Fig. 5E**). The resulting *VLDLR* gene copies overlap with segregating structural variants impacting one or more exons in many duck species (**Fig. 5E inset**). *VLDLR* genes play important roles in the synthesis of yolk protein precursors in ducks^55^, and have recently been shown to facilitate the entry of various alphaviruses into diverse vertebrate species and even in nematodes^56^.

This *VLDLR* expansion may therefore have important implications in ducks’ reproduction as well as viral defense. We observed similar gene expansions accompanied by intraspecific copy number differences in passerine birds, which share a tandem expansion of taste 2 receptor genes (*TAS2R*, 7-22 copies in passerines, 1-9 copies in other birds, **Fig. 5F**)^57,58^. These taste receptors are essential for distinguishing bitter compounds that may be toxic. Extensive variation in their *TAS2R* repertoire may thus have important implications for toxin avoidance in passerine birds. Lastly, GDSL−like Lipase/Acylhydrolase family domains exhibit expanded copy number as well as enrichment for SVs in ray-finned fishes of the clade Otophysi and independently in the Florida worm lizard (*Rhineura floridana*) (**Fig. 5G**). This protein domain is shared by many genes with diverse functions, while the specific genes underlying these expansions remain largely uncharacterized and thus warrant further investigation. Together, SVs exhibit the capacity to contribute to genetic load and adaptive genetic variation with extensive SV enrichment observed across vertebrates in genes involved with immunity, sensory perception, and metabolism.

## Discussion

Here, the recent availability of haplotype-resolved assemblies of many vertebrate species enables us to present a comprehensive characterization of structural and single nucleotide variation across the vertebrate tree of life. We uncover a remarkable shift in the composition of genetic variation: SVs constituted a substantially larger fraction of genetic variation in ancestral vertebrates with their contribution declining sharply in the common ancestor of amniotes and further in mammals, crocodilians, and birds. We find that non-repetitive SVs and TEs contribute independently and asynchronously to these declines. However, the underlying causes of the decline in the contribution of structural variants to genetic diversity are challenging to ascertain. One potential contributor is the transition of amniotes to terrestrial environments, which coincides with a reduction in the horizontal transfer of TEs^41^. Another factor influencing the composition of genetic diversity across taxa is variation in SNV mutation rates. Although we attempted to account for this factor, direct mutation rate measurements are still extremely scarce across vertebrates with only 36 of the species we assessed having known SNV mutation rates, out of which 24 are mammals^10,13^. Thus, further work will be needed to fully disentangle these effects. Taken together, these data highlight the importance of quantifying mutation rates of diverse forms of genetic variation, including both SNVs and more complex SVs, across diverse taxa.

Our finding that the composition of genetic diversity is different across taxa has important implications for vertebrate evolution. Because SVs are more likely to strongly disrupt protein-coding sequences than SNVs, the average effect size of a mutation in fishes and amphibians may be larger, and potentially more deleterious, than in amniotes. This suggests major differences in the distribution of fitness effects, and potentially even the mutational load across vertebrate clades. Fishes and amphibians could potentially experience greater exposure to deleterious mutations within a single generation, which would in turn influence genome and life history trait evolution^59–61^. Conversely, the greater abundance of SVs may also provide increased access to large-effect variants in fishes and amphibians that may be adaptive in novel environments or lead to novel traits^62,63^. Moreover, SVs are known to disrupt genomic stability, drive karyotype evolution, and sometimes contribute to reproductive isolation and speciation^25^. The relatively high abundance of SVs in fishes and amphibians may therefore contribute to their extensive karyotypic diversity, whereas birds, which have the lowest SV content in vertebrates, are characterized by highly conserved karyotypes^64,65^. Finally, although our analyses focus on germline mutation, many SV mutational mechanisms occur somatically as well^66^. Somatic SVs are associated with cancer and other diseases^2^, thus differences in the relative contribution of germline SVs across vertebrate genomes may translate into differences in somatic mutation profiles and age-associated disease susceptibility.

Broadly, our study underscores the extraordinary diversity of SVs as a mutational class. Compared with SNVs, SVs vary across many additional dimensions including size, sequence content, mechanism of formation, and functional consequence^67^. Each of these properties has undergone extensive evolutionary changes across the vertebrate phylogeny, highlighting the highly dynamic nature of SV evolution. One of the core concepts of evolutionary biology is the molecular clock, which relies on the relatively stable accumulation of SNVs across species over time^68^. The rapid turnover and extensive heterogeneity we observe in SVs suggest that they follow fundamentally different mutational and evolutionary dynamics. Unlike SNVs, which are a relatively uniform source of small-scale variation, SVs have the capacity to evolve in bursts and introduce large-scale changes to the genome. As a result, SVs provide a critical substrate for rapid phenotypic evolution and adaptation, enabling punctuated evolutionary changes in contrast to the more gradual dynamics typically associated with SNVs^19^. Together, our results demonstrate that genetic variation across the tree of life cannot be fully understood through SNVs alone, and that SVs represent a major and evolutionarily dynamic component of genetic diversity with broad implications for evolutionary, population, and conservation genetics. The generation of high-quality haplotype-resolved genome assemblies, pioneered by initiatives such as the VGP, CCGP, and CBP, thus represents a paradigm shift, enabling discoveries previously impossible with pseudohaploid reference genomes supplemented by short-read sequencing. Nevertheless, even these resources do not yet fully capture the most extreme forms of structural variation. In particular, very large ampliconic gene arrays remain difficult to assemble, phase, and study because their near-identical repeated copies can exceed the resolving power of standard sequencing and assembly approaches^69^. Likewise, large SVs that span unresolved regions or differ between incompletely assembled haplotypes may remain inaccessible or be represented only partially. Continued improvements in long-read sequencing, assembly, and pangenomic methods will enable population-scale long-read resequencing, near-T2T haplotype assemblies for multiple individuals in the same species, and increasingly accurate SV discovery across diverse taxa^70–72^.

These advances are poised to transform our understanding not only of how biodiversity evolves, but also of how mutational landscapes themselves change across the tree of life.

## Online content

Supplementary figures, tables, and text can be found in Supplementary Online Materials.

## Methods

### Dataset curation

We downloaded the metadata for all vertebrate genome assemblies available on NCBI as of 04/28/25 from the NCBI Genome Dataset page. Among these, we selected the ones that are haplotype-resolved (i.e. a pair of primary and alternate haplotypes for the same biosample) and are at least partly based on PacBio data (i.e. not Illumina- or ONT-only). We required the contig N50 of both haplotypes to be above 100 Mbp. We removed unbalanced pairs of assemblies, where the assembly span of the shorter haplotype is less than 70% that of the longer haplotype. We also removed assemblies from hybrid individuals at the species level. We allowed a maximum of two assembly pairs per species, except for two cases where three were included due to their distinct subspecies status (*Peromyscus maniculatus* and *Chrysemys picta*). When the designation of primary vs. alternate haplotype was not obvious based on the assembly name and assembly type, the more contiguous assembly was chosen as the primary haplotype. This final dataset includes 674 assemblies from 629 vertebrate species. We downloaded the selected assemblies from NCBI using the NCBI datasets tool^73^. We additionally obtained gene annotations for these assemblies when they were available on NCBI or GenomeArk (n=415).

The IUCN conservation status for all species was obtained from the IUCN database using the rredlist R package^74^. A phylogenetic tree scaled to time was obtained from TimeTree5^75^, which includes 632 assemblies from 609 species. An unscaled phylogenetic tree with all 629 species was obtained from the Open Tree of Life^76,77^. The timetree was used throughout the main text when a time-calibrated tree was required for data visualization, phylogenetic correction, or ancestral state reconstruction, resulting in a slightly smaller number of species being included in these analyses. Results from the full unscaled tree were included in some Extended Data Figures and supplementary tables, where branch lengths were obtained using Grafen’s method^78^ when they are needed for phylogenetic correction (e.g. **Extended Data Figure 2**). Major findings presented in this paper were not affected by these different phylogenetic trees.

### Variant calling

First, we split the alternate haplotypes with scaffolds/contigs longer than 270 Mbp into smaller fragments under 200 Mbp to prevent a downstream computational issue. For each sample, we aligned the alternate haplotype to the primary haplotype using minimap2-v2.29^79^ (-a -x asm5 --cs -r2k). We converted the resulting sam files to paf format using the paftools script in minimap2, and used paftools to call SNVs and insertion/deletions of all sizes (call -l 2000 -L 10000). We also generated a bed file specifying the callable regions as determined by paftools using the same parameters. In addition, we sorted the sam files and converted them to bam format using samtools-v1.17^80^, and called SVs using svim-asm-v1.0.3^81^ with the haploid mode. We extracted the SV sequences from variant vcf files using bcftools-v1.9^82^. Paftools does not call duplications and inversions whereas svim-asm does. However, because paftools has a clearer definition of callable regions and insertions/deletions represent the vast majority of all SVs that we identified, we chose to present results from paftools throughout the paper unless otherwise noted.

### Statistical analysis of SV and SNV heterozygosity

SNV and SV heterozygosities were computed as the number of variants divided by the length of the callable region. To compare the differences in heterozygosity between major vertebrate clades and IUCN conservation categories, we took a phylogenetic generalized least squares (PGLS) approach using Pagel’s λ^83^ to account for correlation structures along the phylogeny and included clades, IUCN category, sequencing technology (i.e. PacBio CLR vs. CCS) as explanatory variables. We followed the PGLS with Tukey’s all-pair comparisons to determine the statistical significance of differences among groups. The relationship between H_SV_ and H_SNV_ was similarly evaluated using PGLS while taking sequencing technology into account. Lastly, we performed ancestral state reconstruction of the SV-to-SNV ratio using a maximum likelihood approach, excluding two outlier assemblies with extremely high or low ratios for clearer visualization. All statistical analyses were performed in R using packages nlme-v3.1^84^, ape-v5.8^85^, multcomp-v1.4^85,86^, phytools-v2.5^87^ and visualized with ggplot2-v4.0.2^88^ and ggtree-v4.1.1^89^.

### Repeat annotation

To annotate different types of repetitive elements in SVs and TEs, we first generated species-specific repeat libraries by combining de-novo libraries with curated public databases^48^. We generated de-novo repeat libraries using RepeatModeler-v2.0.5^90^ (-LTRStruct -engine rmblast). RepeatModeler could not be run to completion for a few species despite multiple attempts, so we downloaded RepeatModeler libraries for the same species from the UCSC genome browser^91^ when they were available, and incomplete libraries when they were not. We obtained curated ancestral and descendant repeats from the Dfam database (v3.9)^92^ for each species using the famdb.py script in RepeatMasker-v4.1.7^93^ (famdb.py families --ancestors --descendants --include-class-in-name --curated). We used an iterative method that started from the species name and proceeded upward in taxonomic order up to infraclass in order to obtain at least 50 curated repeat sequences for each species. We reduced the redundancy in the de-novo and Dfam libraries by running cd-hit-est-v4.8.1^94^ (-d 0 -aS 0.8 -aL 0.8 -c 0.8 -G 0 -g 1 -b 500) with each library separately, giving priority to named vs. unknown repeats. We combined the de-novo RepeatModeler library with the curated Dfam library and used cd-hit-est to perform another round of redundancy reduction. We ran RepeatMasker (-engine rmblast -gff -lib) on the genomic and SV sequences using these species-specific repeat libraries. We noticed that the Dfam library alone outperforms the combined library for primates due to their high level of representation in Dfam. We therefore chose to use the dfam library without RepeatModeler for primates. An SV is considered to be of a certain repeat type if more than 70% of its sequence is annotated by RepeatMasker and if the top repeat type occupies more than 70% of the annotated part of the sequence. Otherwise it is labeled as a “non-repeat”.

Unclassified repeats are treated as TEs in **Fig. 2** because they are likely TEs yet to be curated in Dfam. For TE-specific analyses (**Fig. 3**), a total of 130 TE families were identified, and 39 most abundant ones among all species were analyzed separately. Others were grouped together into combined classes including SINE/other, LINE/other, etc. We computed the Shannon Diversity Index (H’)^95^ for TEs in each species. We performed this computation through scikit-bop^96^. To identify differences in the TE composition in SVs vs. genomes, we performed a Fisher’s exact test comparing the proportion of each classified TE family among all TEs in each species’s SV sequences and genome by TE count. A Bonferroni correction was performed on all tests to account for multiple testing. Note that in this analysis, statistically significant turnovers of a TE family in multiple species do not imply multiple independent events as they may have occurred in a single common ancestor.

### Genomic substrate of SVs and enrichment of k-mers and non-canonical DNA structure

To investigate the genomic substrate of SVs, we trained a random forest classifier using scikit-learn-v1.6.1^97^ to predict SV occurrence from nearby genomic features. We extracted 400-bp windows flanking each SV (200 bp upstream and 200 bp downstream), excluding the SV sequence itself, to form a positive set. An equal number of 400-bp windows that do not contain SVs were randomly sampled from the genome to form a negative set.

We randomly sampled 80% of these for model training and used the other 20% for testing. We obtained genomic features within each 400-bp window including the number of SNVs, small insertions/deletions <50 bp, inversions, distance to different repetitive elements, and GC content, and used them as predictors for the random forest model. For a subset of species that have gene annotation available, we additionally included distance to the nearest exon in the model, but this did not make a substantial difference to the predictive power likely due to redundancies with other features such as the number of SNVs. We thus chose not to present the result with gene annotations in the main text. We used 1000 trees in each random forest with balanced class weights, maximum number of features when splitting as the square root of the total number of features, and default bootstrapping. All models were evaluated on the held-out test set. To evaluate feature importance, we computed permutation importance, which measures the decrease in model performance when individual features are permuted. Permutation importance was calculated on the test set to provide an out-of-sample estimate, with 5 repetitions. Correlations between features and SV presence/absence were calculated using the test set as well.

To characterize the sequence content around SVs, we counted the 5-mers within all 400-bp windows used for the random forest model. Furthermore, we searched for motifs associated with G-quadruplex (G4) and Z-DNA structures within these 400-bp windows, following the definition by Makova and Weissensteiner^47^. Enrichment was computed using all SVs against the balanced negative set.

### Sequence homology across SV breakpoints

To characterize patterns of homology across SV breakpoints, we roughly followed the workflow established by Schloissnig et al.^29^ Briefly, for each SV, we extracted sequences immediately upstream and downstream of the SV breakpoint in the reference genome with length equal to half of the SV length, and concatenated them with the first and second halves of the SV sequence, respectively. We then blasted these pairs of sequences against each other using blastn-v2.17.0^98^ (-perc_identity 80 -word_size 5) to find regions of homology between them. We only considered homologies that were in the direct orientation, that crossed the SV breakpoint with a 10-bp cushion, and that were located around the same relative position in either flank. If multiple homology alignments met these criteria for a single SV, we retained the longest spanning one. We grouped all SVs into four categories based on their homology length in comparison to the SV length: “non-homology-associated” if no homology was found, “microhomology-associated” if the homology length was <30 bp and <40% of the SV length, “homology-associated” if the homology length was >=30 bp and <40% of the SV length, and “tandem events” if the homology length was >=40% of the SV length.

To identify the mechanisms through which homologies can be linked to the SVs, we further annotated the repeat type of the homology sequences with RepeatMasker in the same method as for SV sequences.

Tandem events were classified as “TE/satellites-asscociated tandem events” if their homology sequences were annotated as TEs or satellites (including unclassified repeats since most of them are likely unclassified TEs or satellites). Microhomology-associated events were classified as “TE-associated TSDs” if the SVs were annotated as TEs (including unclassified repeats). Homology-associated events were classified as “LTR-associated long terminal repeats” if the SVs were annotated as LTRs. Homology-associated events were classified as “TE-mediated insertion/deletion” if their homology sequences were annotated as TEs (excluding unclassified repeats to be more conservative) and the SVs were annotated as non-repeats.

### Variant effect

We used bedtools-v2.30.0^99^ to count the overlap between genetic variants and different genomic features in the reference genome, including intergenic, gene, intron, exon, UTR, and CDS sequences. Enrichment ratios were computed as the observed divided by the expected length of overlap. We used Ensembl VEP-v113.3^52^ to evaluate the potential functional impact of each genetic variant and computed the proportion of variants with predicted “HIGH” impact on at least one protein-coding gene for each species. To quantify the enrichment of genetic variants in different protein domains, we first extracted the longest protein sequence per gene in each assembly using gffread-v0.12.7^100^. We used interproscan-v5.59-91.0^53,101^ to annotate functional domains and ontologies in these protein sequences. We converted the domain annotations from the protein space back to the genomic space using custom scripts, and assessed overlap between genetic variants and protein domains using bedtools. Enrichment ratios and p-values were obtained for deletions from 10000 permutations using GAT-v1.3.6^102^. For SNVs and insertions, since one base pair is affected at a time, enrichment ratios and p-values could simply and accurately be obtained from a binomial process. Regions of the genome that could not be called by paftools were excluded from all enrichment analyses. Multiple testing correction was not performed on these p-values because our selection of domains of interest does not rely on a single statistically significant test but on enrichment patterns that are consistent across species or that exhibit strong phylogenetic patterns. Synteny plots in select species and genes were constructed in R using the package SVbyEye-v0.99.0^103^.

## Data Availability

All data used in this paper are publically available and described in the “dataset curation” section of the methods. Assembly accession numbers can be found in **Table S1**. All species illustrations were either obtained from the public domain or created for this paper.

## Code Availability

All code used in this paper can be found in the following GitHub repository https://github.com/sudmantlab/sv_diversity and is archived on Zenodo (doi:10.5281/zenodo.20804765).

## Acknowledgements

We thank the Vertebrate Genomes Project (VGP), particularly Erich Jarvis and Giulio Formenti for their leadership, and Heng Li, Aryn Wilder, and Amanda Gardiner for valuable discussions and feedback. We are also grateful to members of the VGP transposable element (TE) working group, particularly Richard Durbin, Clément Goubert, Robert Hubley, Hiram Clawson, and Shujun Ou, for their insights and guidance on TE analyses. We further acknowledge the substantial contributions of other sequencing consortia and institutions, including the California Conservation Genomics Project, Canadian Biogenome Project, Bat1K, and the Wellcome Sanger Institute, whose efforts helped generate the datasets used in this study. We are deeply appreciative of the insightful discussions, encouragement, and constructive feedback provided by members of the Sudmant lab throughout this project, especially Caitlin Connelly for her careful review of the manuscript. Finally, we thank Philippa Steinberg for creating the beautiful species illustrations featured throughout the manuscript.

This work was supported by NIH National Institute of General Medicine award R35GM142916 to PHS. This manuscript is the result of funding in whole or in part by the National Institutes of Health (NIH). It is subject to the NIH Public Access Policy. Through acceptance of this federal funding, NIH has been given a right to make this manuscript publicly available in PubMed Central upon the Official Date of Publication, as defined by NIH. The content is solely the responsibility of the authors and does not necessarily represent the official views of the National Institutes of Health.

## Contributions

Conceptualization: RNL, PHS

Data curation and formal analysis: RNL, DL, MD, LG, PHS

Writing – original draft: RNL, PHS

Writing – review & editing: RNL, DL, LG, GLO, PHS

Resources: Vertebrate Genomes Project Consortium

Funding acquisition: PHS

Supervision: NI, PHS

## Extended Data Figures

**Extended Data Figure 1.**
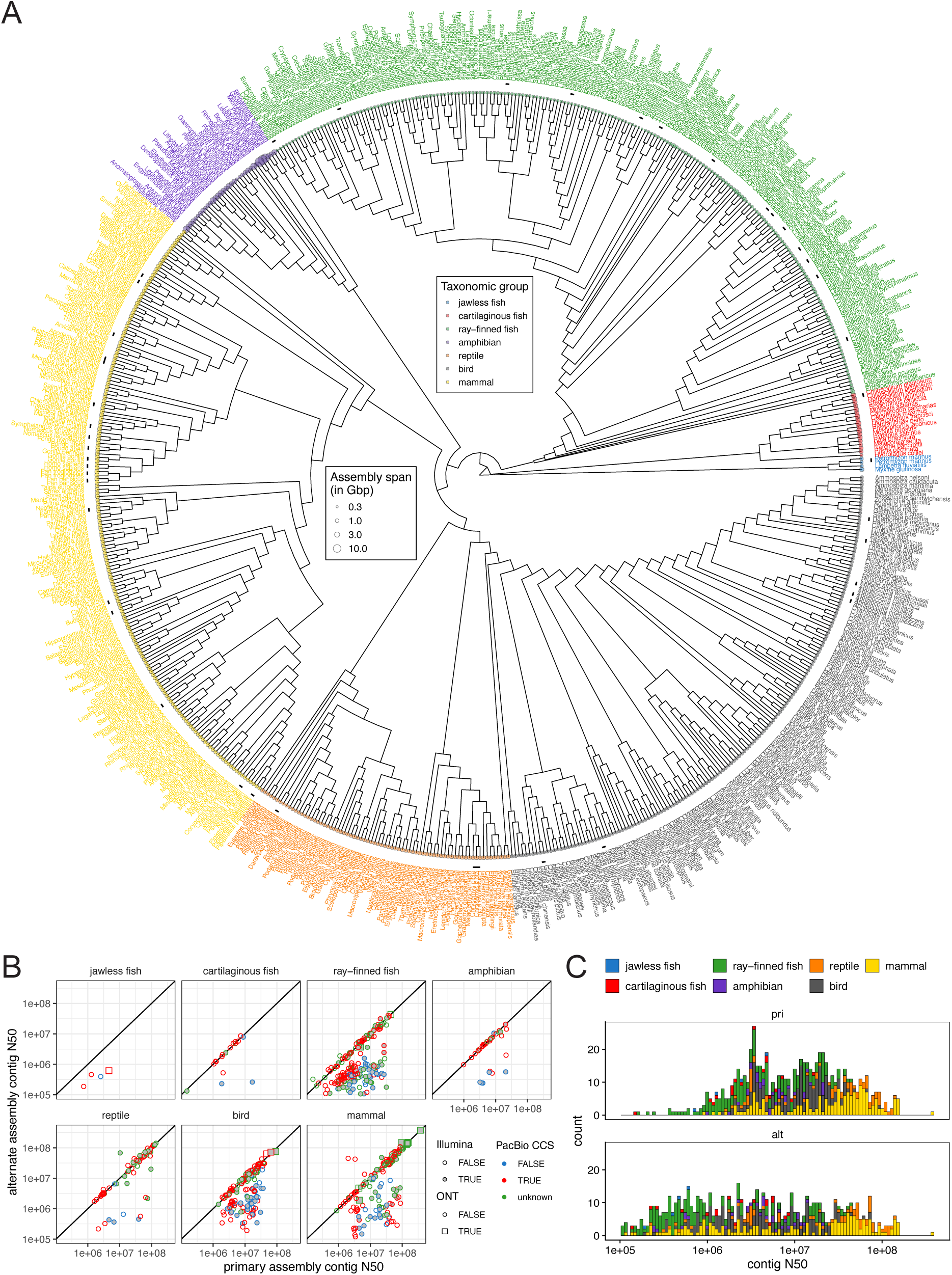
An overview of the dataset. **A)** An unscaled phylogenetic tree of all 674 genome assemblies included in this paper. Species names are shown at the tip of the tree. Colors indicate major vertebrate clades. Multiple assemblies of the same species are indicated with short lines next to their tips. The lengths of primary assemblies are indicated by the size of circles at the tip of the tree. **B)** The relationship between the contig N50 values of the primary and alternate assemblies, grouped by clade. A 1-to-1 line is drawn in each plot. Colors of the points correspond to the type of PacBio sequencing technology used. Fills of the points indicate whether Illumina short-read data were included in the assembly. Shapes of the points indicate whether ONT data were included in the assembly. **C)** The distribution of contig N50 values of primary and alternate assemblies, colored by clade.

**Extended Data Figure 2.**
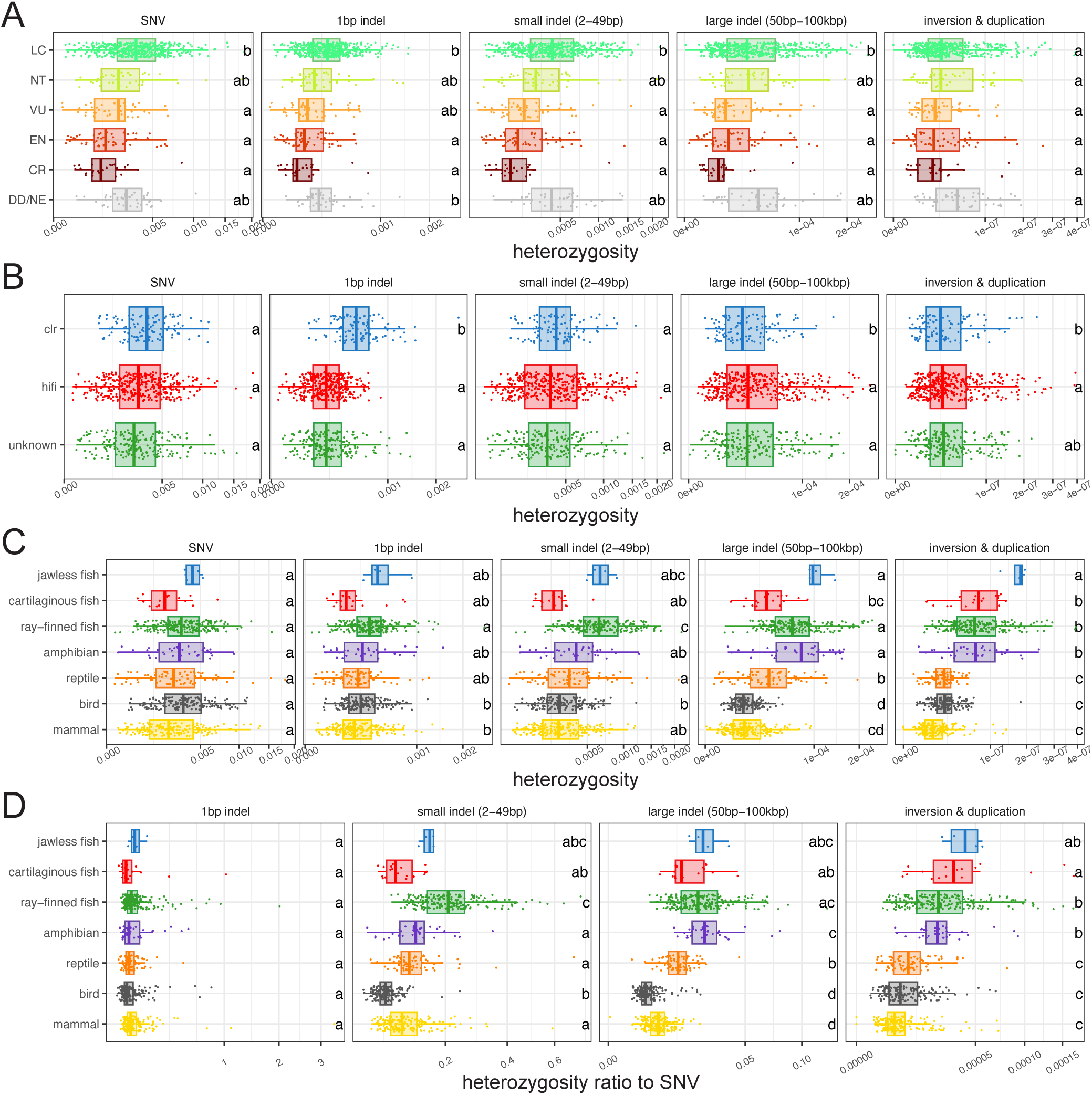
The distribution of different classes of genetic variation across 674 assemblies. A-C) The distribution of SNV, 1-bp indel, small indel (2-49 bp), large indel (50 bp-100k bp), and inversion/duplication heterozygosity grouped by IUCN conservation category (**A**), PacBio sequencing technology (**B**), and clade (**C**). **D)** The distribution of the ratios of 1-bp indel, small indel (2-49 bp), large indel (50 bp-100 kbp), and inversion/duplication to SNV heterozygosity. Different letters on the right indicate statistically significant differences between categories (Bonferroni-adjusted p-value threshold of 0.05, phylogenetic generalized least squares followed by Tukey’s test). This figure includes all 674 genome assemblies analyzed in this paper. Phylogenetic correction was done using an unscaled tree with Grafen’s branch lengths. Fig. 1 **D-I** shows the subset of 632 assemblies that can be placed on a time-calibrated tree.

**Extended Data Figure 3.**
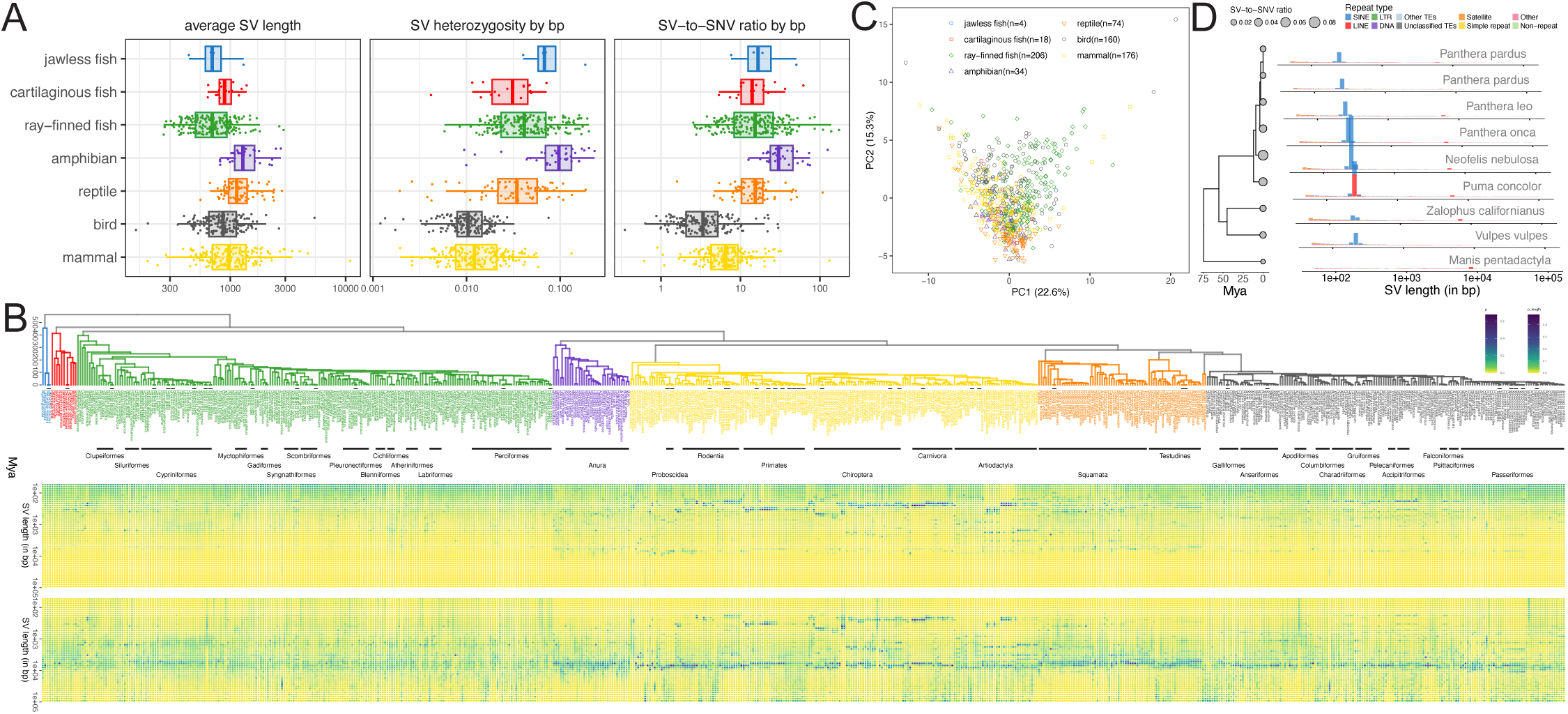
The distribution of SV length across species. **A)** The distribution of average SV length, SV heterozygosity measured by base pairs impacted, and SV-to-SNV ratio measured by base pairs impacted, grouped by clade. **B)** A heatmap of SV length distribution mapped onto a time-calibrated phylogenetic tree. SVs are grouped into bins of different sizes on the y-axis, and colors indicate the proportion of SVs of a given length bin out of all SVs. The proportion is measured by count in the top panel, and length in the bottom panel. **C)** A PCA of length distribution across species colored by clade. **D)** SV composition and length distribution in nine carnivores. The size of the circles at the tips of the phylogeny indicates the SV-to-SNV ratio in each assembly. To their right, the histograms show the length distribution of SVs normalized by SNVs, colored by SV type. Note that the x-axis in each example is slightly offset to reduce overlap between plots.

**Extended Data Figure 4.**
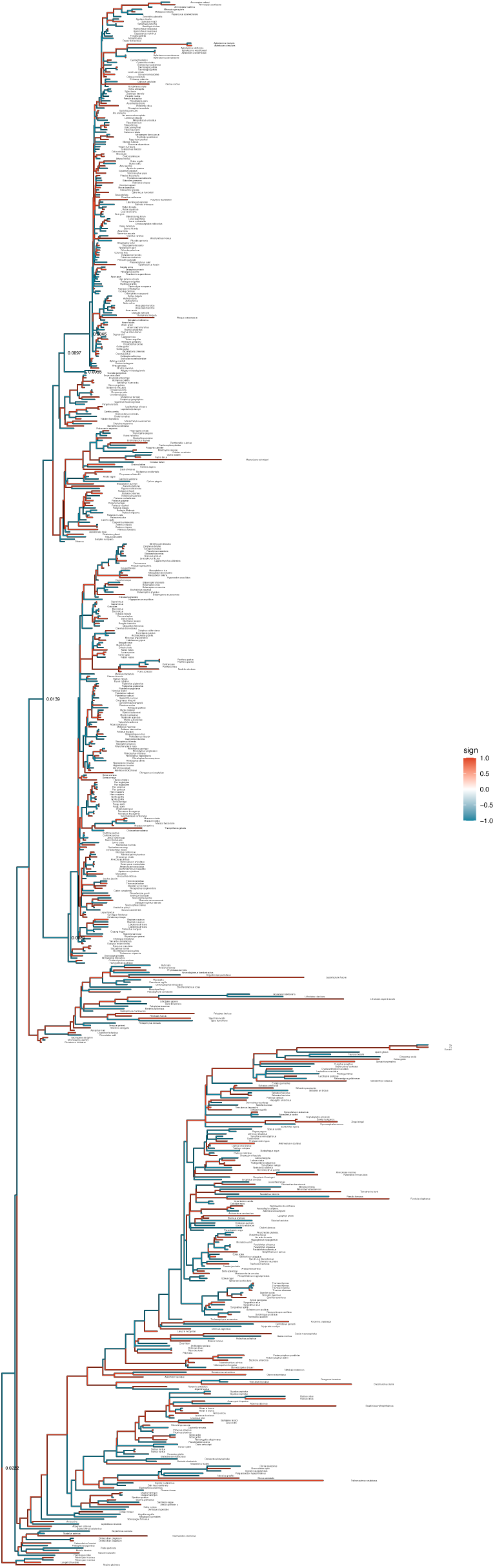
Ancestral state reconstruction of SV-to-SNV ratio. Branch lengths correspond to the amount of change in SV-to-SNV ratio along each branch, whereas the colors indicate the direction of change (red: increase, blue: decrease). SV-to-SNV ratios at certain ancestral nodes are annotated on the tree.

**Extended Data Figure 5.**
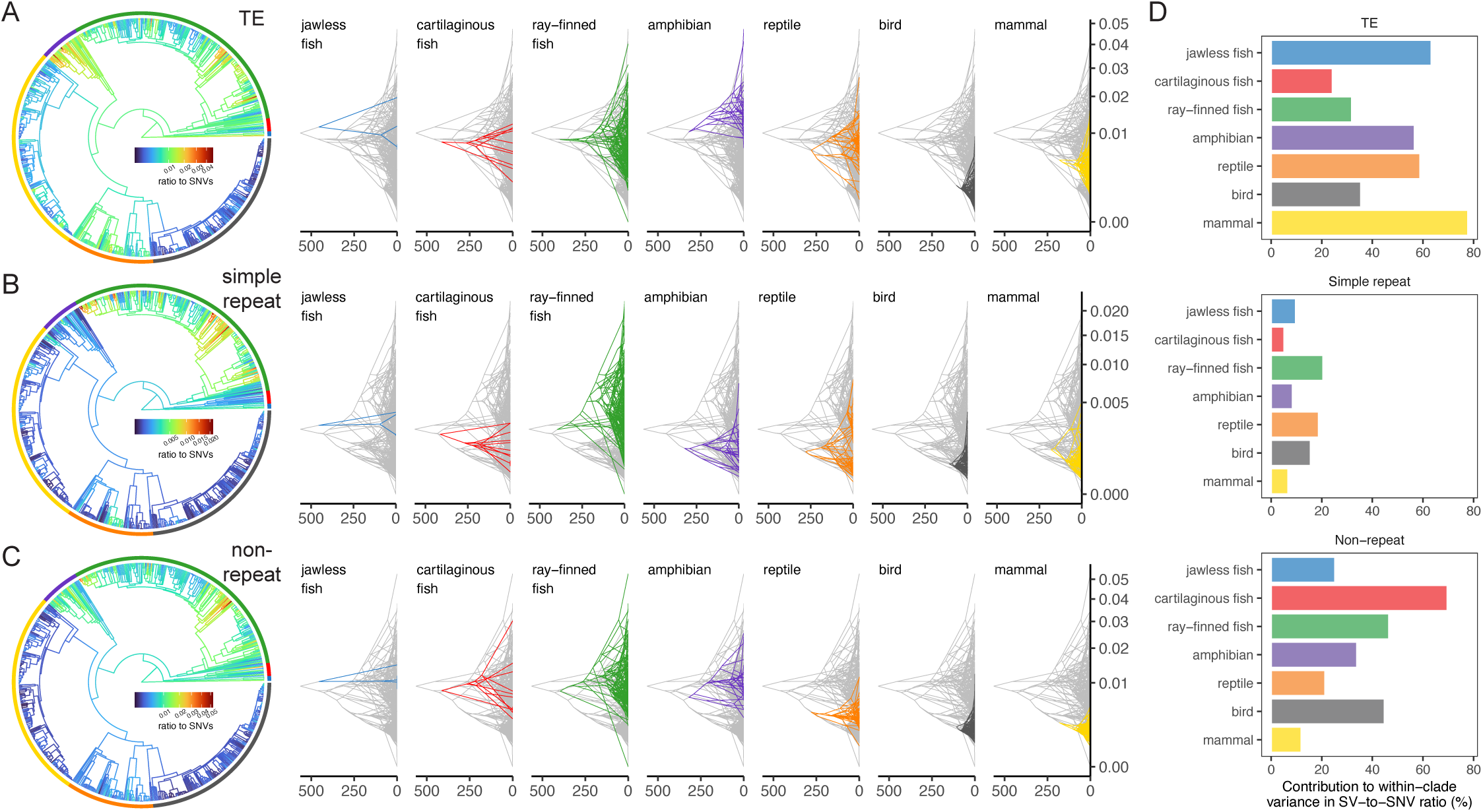
Contributions to the SV-to-SNV ratio by different types of SVs. A-C) Ancestral state reconstruction of SV-to-SNV ratio broken down by SV types, shown as circular trees on the left and slanted trees on the right. In circular trees, colors at the tips indicate the clade, and colors on the tree indicate ancestral trait values. **D)** The contribution to within-clade variance in SV-to-SNV ratio by different SV types.

**Extended Data Figure 6.**
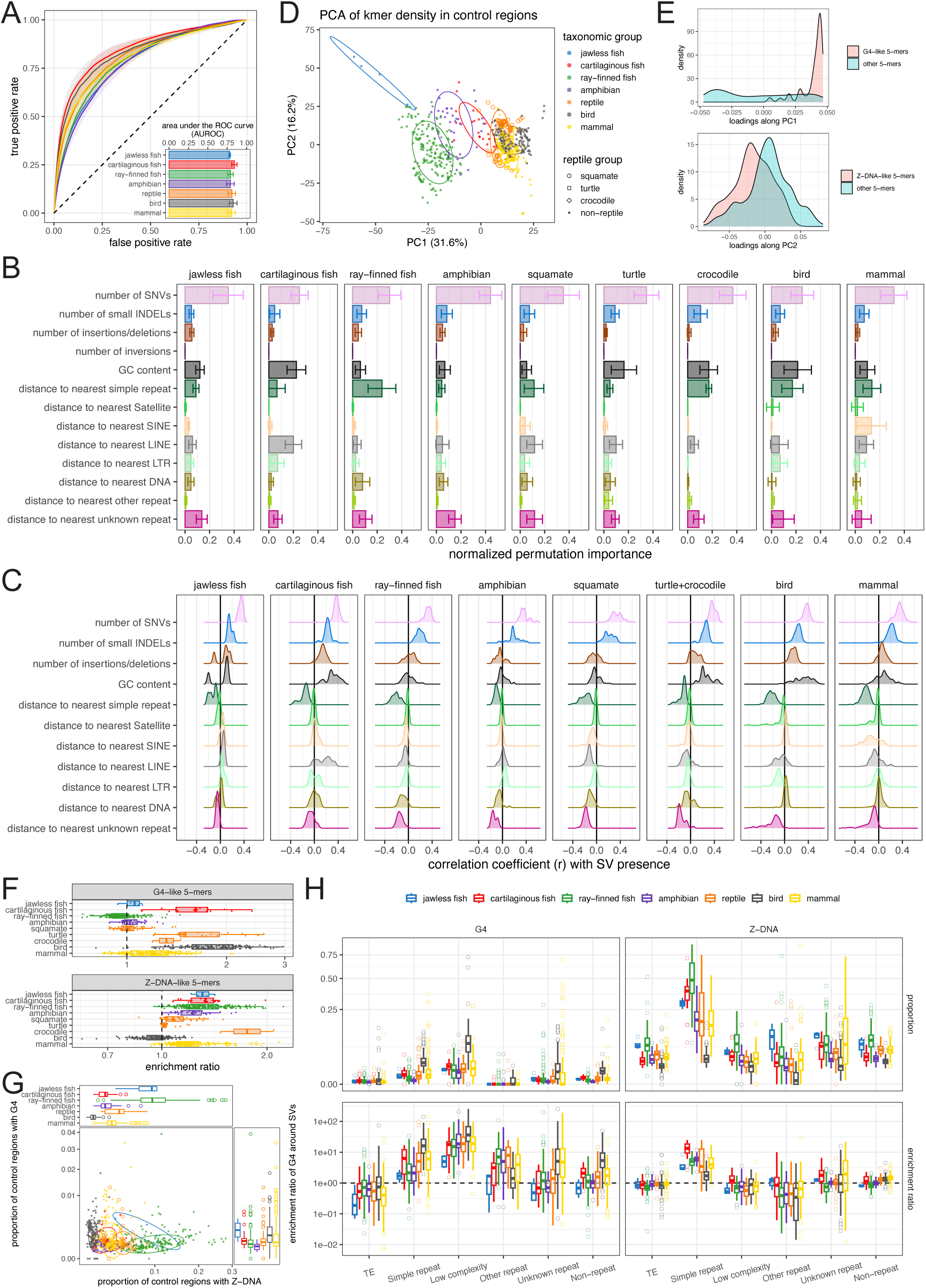
Genomic substrates of SVs. **A)** Receiver Operating Characteristic (ROC) curves of random forest models in predicting SV occurrence given genomic annotations in a 400-bp window. Colored lines show the median in each clade, whereas shades along the lines indicate the interquantile range. **B)** The normalized permutation importance of genomic features in predicting SV presence grouped by clade. Bars indicate the median value in each clade and error bars indicate the standard deviation. **C)** The distribution of correlation coefficients between different genomic features and SV presence/absence, grouped by clade. Note that repetitive elements are coded by the distance to them, so negative correlations suggest greater proximity between repetitive elements and SVs. **D)** PCA of 5-mer densities in the genomic background across species, colored by clade. Different shapes further distinguish three clades within reptiles. **E)** The distribution of PC loadings along PC1 (top) and PC2 (bottom) in the 5-mer enrichment PCA shown in Fig. 4B. 5-mers are divided into G4-like 5-mers vs. others on top, and Z-DNA-like 5-mers vs. others at the bottom. **F)** Enrichment of G4-like (top) and Z-DNA-like 5-mers around SV breakpoints, grouped by clade. **G)** The proportion of 400-bp windows with G4 and Z-DNA motifs in genomic sequences, colored by clade. Figure legend is the same as in **D)**. **H)** Enrichment of G4 and Z-DNA motifs around SV breakpoints, broken down by different SV types on the x-axis, colored by clade.

**Extended Data Figure 7.**
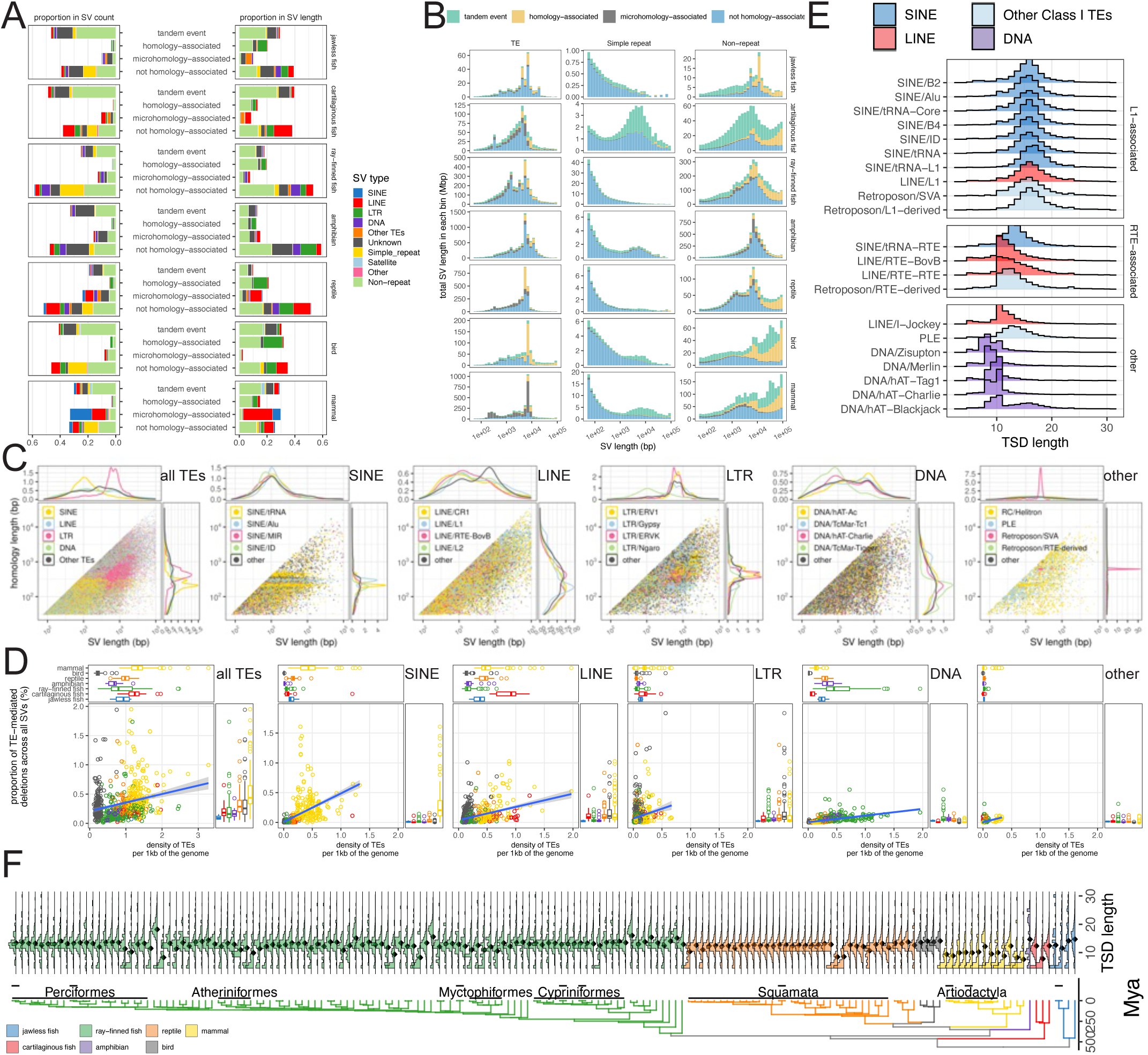
Patterns of homology across SV breakpoints. **A)** The proportion of SVs with each homology category in each clade, quantified by the number of SVs (left) and the number of base pairs impacted (right). Colors correspond to SV type. **B)** The length distribution of SVs grouped by SV type and clade, and colored by their homology category. Y-axis is the total length of SVs falling into a given length bin. **C)** The distribution of SV length and homology length in insertions/deletions mediated by all TEs (panel 1) and TEs of different orders (SINE, LINE, LTR, DNA, others, panels 2-6). Points are colored by TE type. Marginal plots show the density distribution along each axis. **D)** The relationship between the proportion of TE-mediated insertions/deletions (y-axis) and TE abundance (quantified as the density of classified TEs in the genome assembly, x-axis), colored by clade and grouped by TE order. **E)** The distribution of target site duplication (TSD) length of different TE families, grouped by the types of endonuclease that they rely on. **F)** The evolution of TSD length associated with LINE/RTE-BovB endonuclease along the vertebrate phylogeny.

**Extended Data Figure 8.**
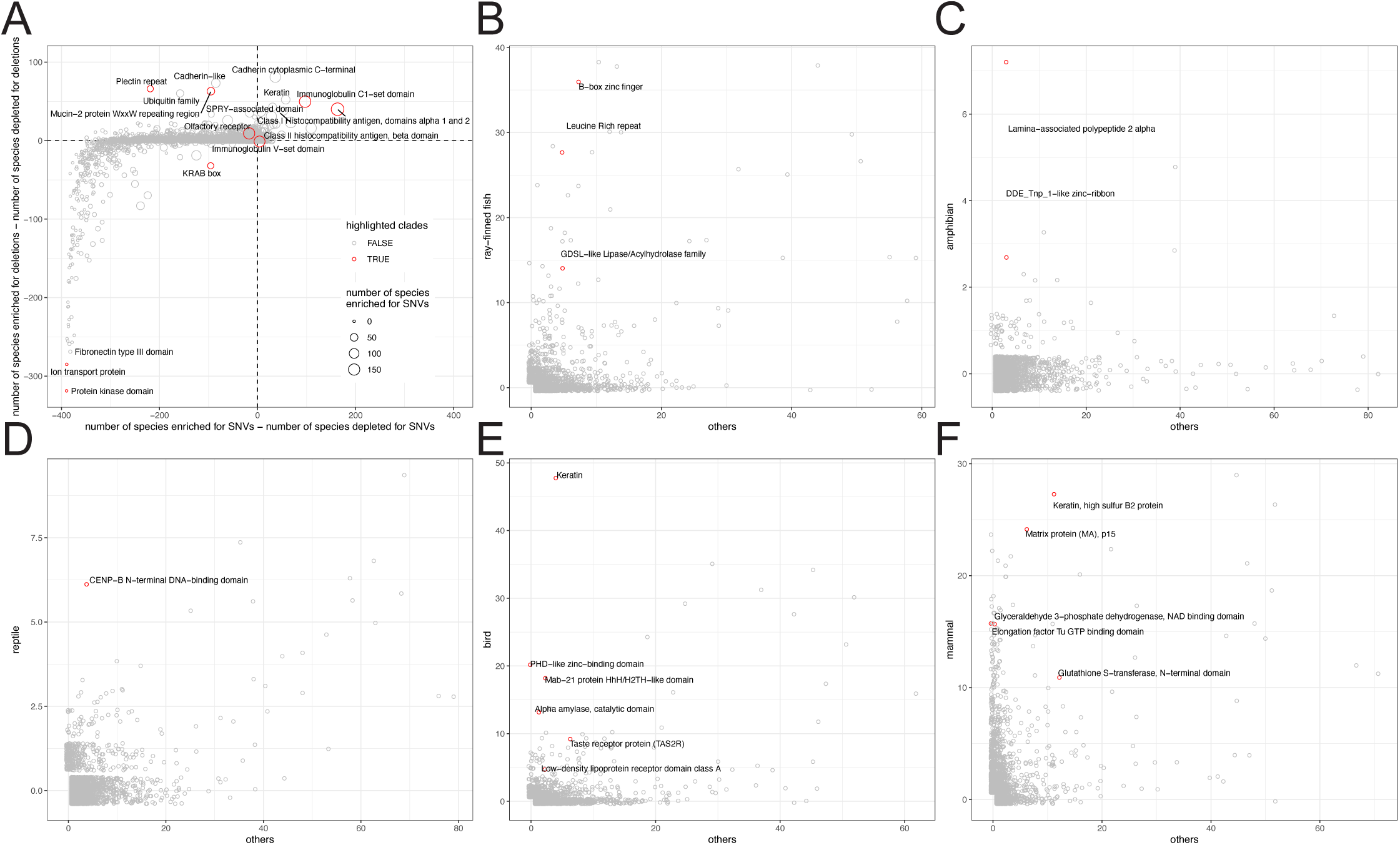
The enrichment and depletion of SVs and SNVs in different protein domains. **A)** Each point is a protein domain. X-axis shows the number of species that are significantly enriched for SNV subtracted by the number of species that are significantly depleted for SNVs per protein domain (p-value threshold of 0.05 without multiple testing correction). Y-axis is the same but for SVs. The size of points corresponds to the number of species that are enriched for SNVs. Select outliers are labeled and examples highlighted in Fig. 5D are colored in red. **B-F)** Domains exhibiting clade-specific patterns of SV enrichment. Each point is a protein domain. Y-axis shows the number of species in each focal clade that are significantly enriched for SVs. X-axis shows the number of species in all other clades that are significantly enriched for SVs. Examples highlighted in Fig. 5D are labeled and colored in red.

