## Supplementary Notes, Supplementary Figures 1-6 for "The evolution of structural variation across 500 million years of vertebrate evolution"

#### Relevance of haplotype-resolved reference genomes in biodiversity conservation

Past research has suggested that genetic diversity is reflective and predictive of a species’ conservation status and extinction risk^1,2^, but they have been limited by the small number of species (mostly mammals) where polymorphism data are available. We therefore examined the relationship between H_SNV_ and H_SV_ with the conservation status established by the IUCN Red List. Threatened and near threatened species tend to have significantly lower heterozygosity in both their SNVs and SVs than species of least concern (phylogenetic generalized least squares followed by Tukey's test, statistical significance with p-values threshold of 0.05 shown in **Fig. 1E**). For example, among the species with the lowest heterozygosity in our dataset are the Vulnerable (VU) Suwannee snapping turtle (*Macrochelys suwanniensis*, H_SNV_=3.87x10^-5^, H_SV_=6.98x10^-7^), the Endangered (EN) Tasmanian devil (*Sarcophilus harrisii*, H_SNV_=1.04x10^-4^, H_SV_=1.27x10^-6^), and the Critically Endangered (CR) Rice's whale (*Balaenoptera ricei*, H_SNV_=2.22x10^-4^, H_SV_=2.50x10^-6^). In contrast, the species with the highest H_SNV_ include the European nightjar (*Caprimulgus europaeus*, H_SNV_=1.1x10^-2^,H_SV_=2.3x10^-5^), the short-beaked echidna (*Tachyglossus aculeatus*, H_SNV_=1.1x10^-2^,H_SV_=6x10^-5^), the Atlantic silverside (*Menidia menidia*, H_SNV_=9.8x10^-3^,H_SV_=4.4x10^-5^), etc., all of which are species of Least Concern (LC) with broad geographic distributions. Strikingly, the top seven species with the highest H_SV_ are all highly abundant deep sea fishes of orders Myctophiformes, Trachichthyiformes, and Beryciformes (H_SV_>2x10^-4^).

However, there are some notable exceptions. For example, the olive python (*Liasis olivaceus*) is a species of Least Concern, but it is among the species with the lowest H_SNV_ (7.42x10^-5^). The individual that was sequenced, however, is zoo animal that was likely bred in captivity and thus may not be representative of populations in the wild. Other initially unexpected results reflect complexities in a species’ demographic history and ecology. The European eel (*Anguilla anguilla*) has a broad geographic range and was once highly abundant. Human activities in the past decades have drastically reduced their population by more than 90%, and it was declared as Critically Endangered as a result^3^. Its contemporary population size, however, is likely much higher than most other Critically Endangered species, and this recent population contraction has not yet had enough time to impact its genetic diversity. Therefore, its H_SNV_ is at 8.37x10^-3^, among the highest in our dataset. On the other end of the spectrum, the European badger (*Meles meles*) is of Least Concern with an unexpected low H_SNV_ (3.36x10^-4^), but this can be explained by their strong population structure consisting of some small and isolated populations, making them susceptible to inbreeding^4,5^.

Despite these exceptions, broad patterns shown in our data suggest that although genetic diversity is not currently considered in IUCN’s designation of conservation categories, threatened and near threatened species tend to harbor lower genetic diversity because of their lower population sizes. Indeed, with some simple mutation rate and generation time estimates, we separately performed a pairwise sequential Markovian coalescent (PSMC) analysis^6^ and showed that threatened and near-threatened species tend to have smaller effective population size (N_e_) historically^7^. Furthermore, Critically Endangered (CR) species saw sharp declines in more recent times, likely driven by inbreeding. Therefore, these haplotype-resolved assemblies prove to be a powerful tool for conservation especially for species that are hard to sample and survey, as long as cautions are taken in their interpretation.

#### Characterizing the length distribution of SVs

Our categorization of SVs based on their repeat content rely on automated repeat annotation, which is notoriously challenging for non-model species. As a complementary analysis, we used the length distribution of SVs across species agnostic of repeat annotations to confirm these findings (**Extended Data Fig. 3B**). In general, most species tend to have more short SVs compared to longer ones. However, in most mammals, peaks in SV length distributions are common and show strong phylogenetic patterns, indicating strong TE activities and frequent turnovers in mammals. This pattern is also true for some bird species, and for amphibians and reptiles to a lesser extent, consistent with the lower TE activities in birds (**Fig. 2B**) and the more diverse TE landscape in amphibians and reptiles (**FIg. 3C**). In contrast, the general lack of peaks and the lower average lengths of SVs in fishes (**Extended Data Fig. 3A**) are reflective of higher contributions of simple repeat and non-repeat in these clades (**Fig. 2B**), as well as more even contributions from a highly diverse repertoire of TEs (**FIg. 3C**). Therefore, our main conclusions should be generally robust to uncertainties in repeat annotations.

#### References

#### 1. Jeon, J. Y. *et al.* Genomic Diversity as a Key Conservation Criterion: Proof-of-Concept From Mammalian Whole-Genome Resequencing Data. *Evolutionary Applications* **17**, e70000 (2024).

#### 2. Wilder, A. P. *et al.* The contribution of historical processes to contemporary extinction risk in placental mammals. *Science* **380**, eabn5856 (2023).

#### 3. IUCN. Anguilla anguilla: Pike, C., crook, V. & gollock, M. *IUCN Red List of Threatened Species* IUCN https://doi.org/10.2305/iucn.uk.2020-2.rlts.t60344a152845178.en (2018).

#### 4. Annavi, G. *et al.* Heterozygosity-fitness correlations in a wild mammal population: accounting for parental and environmental effects. *Ecol Evol* **4**, 2594–2609 (2014).

#### 5. Pope, L. C., Domingo-Roura, X., Erven, K. & Burke, T. Isolation by distance and gene flow in the Eurasian badger (Meles meles) at both a local and broad scale. *Mol Ecol* **15**, 371–386 (2006).

#### 6. Li, H. & Durbin, R. Inference of human population history from individual whole-genome sequences. *Nature* **475**, 493–496 (2011).

7. Formenti. G. *et al.* The Vertebrate Genomes Project Phase I: A global reference genome resource. *bioRxiv* (2026). doi:10.64898/2026.06.24.732306.

### Supplementary Figures

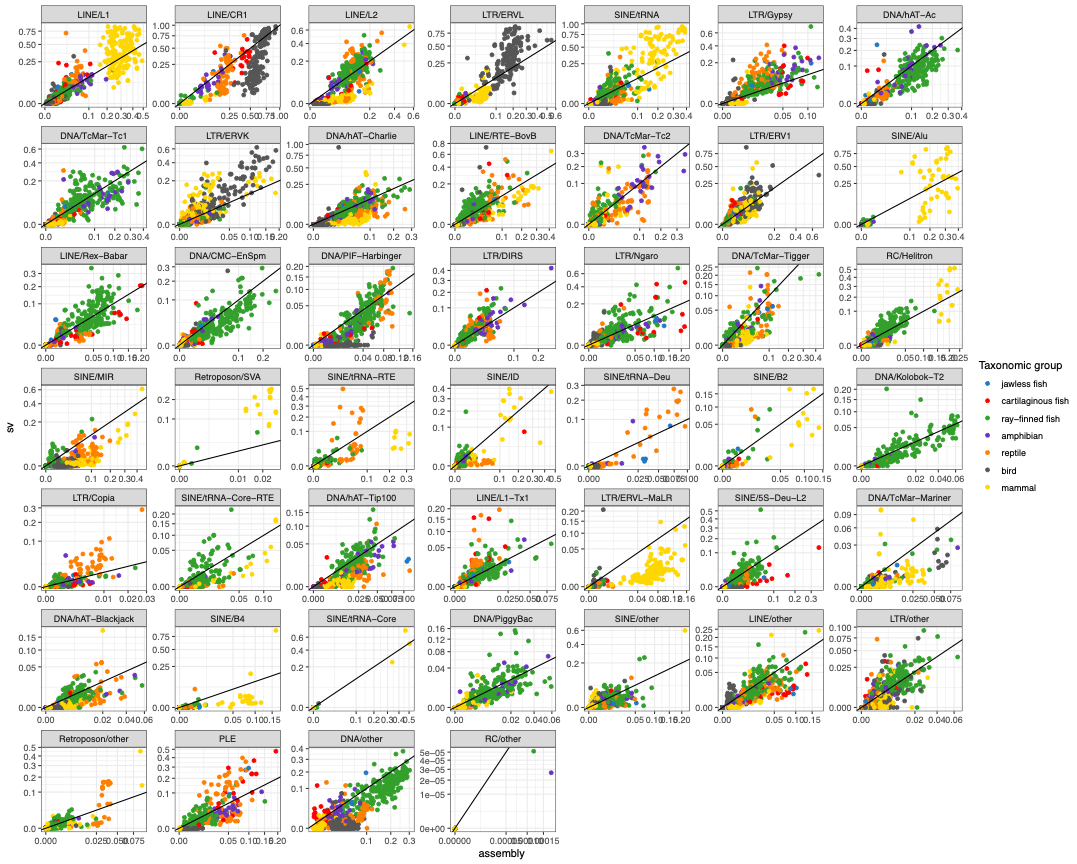

**Figure S1 -** The abundance of all major TE families in SVs (y-axis) and in genome assemblies (x-axis) across major vertebrate clades. TE abundance is quantified by their proportion among all classified TEs. Assemblies are colored by major clades. A one-to-one line is drawn to indicate the expected distribution if TE composition is identical between SVs and assemblies.

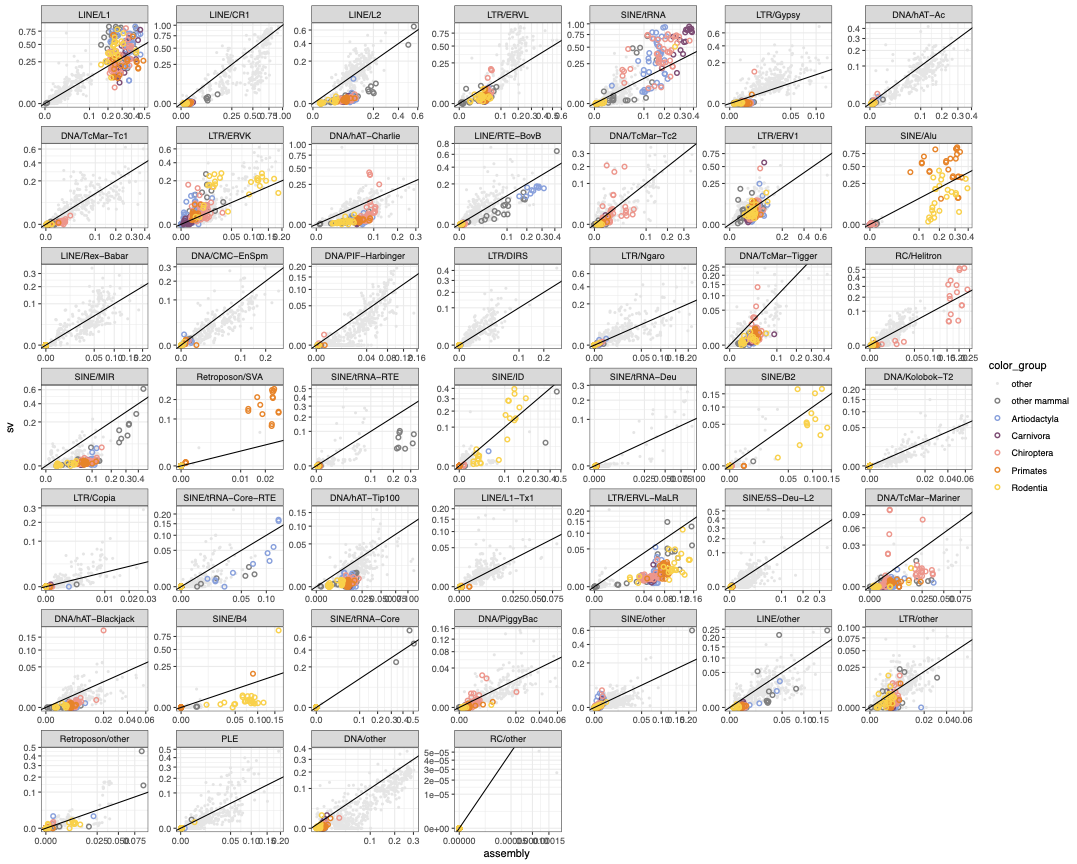

**Figure S2 -** The abundance of all major TE families in SVs (y-axis) and in genome assemblies (x-axis) across mammals. TE abundance is quantified by their proportion among all classified TEs. Assemblies are colored by major mammalian orders. A one-to-one line is drawn to indicate the expected distribution if TE composition is identical between SVs and assemblies.

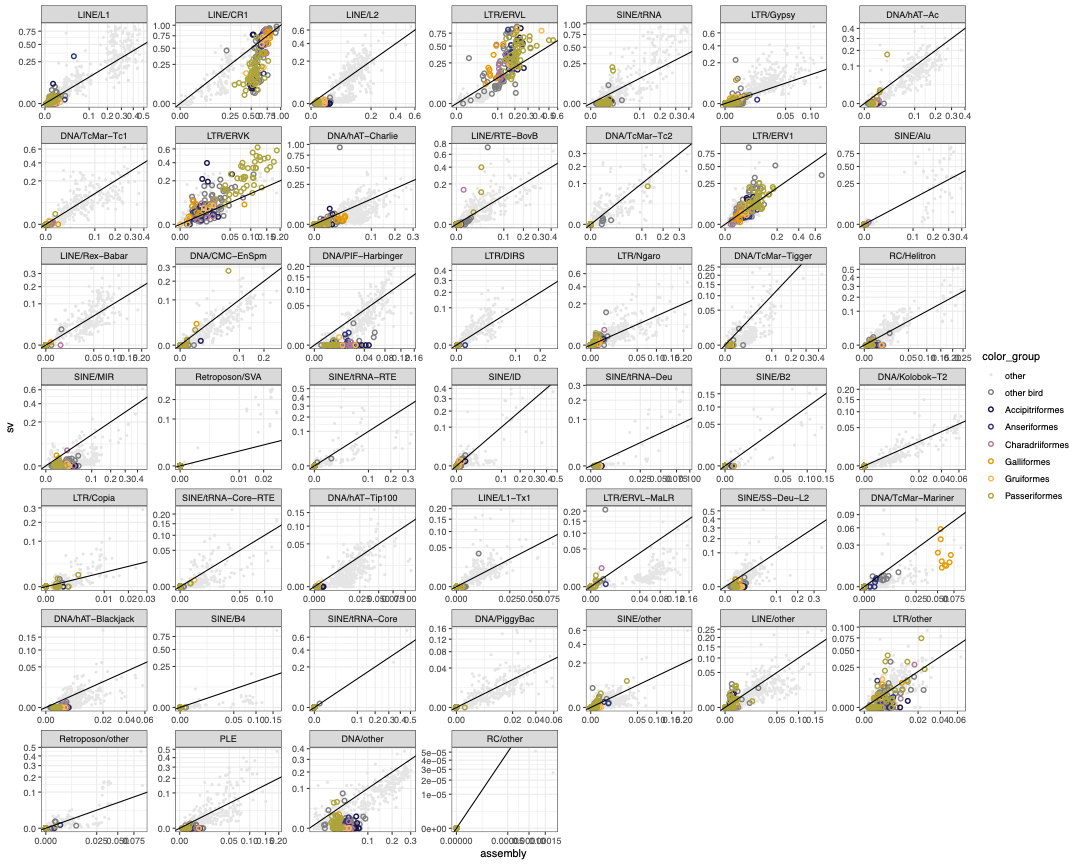

**Figure S3 -** The abundance of all major TE families in SVs (y-axis) and in genome assemblies (x-axis) across birds. TE abundance is quantified by their proportion among all classified TEs. Assemblies are colored by major bird orders. A one-to-one line is drawn to indicate the expected distribution if TE composition is identical between SVs and assemblies.

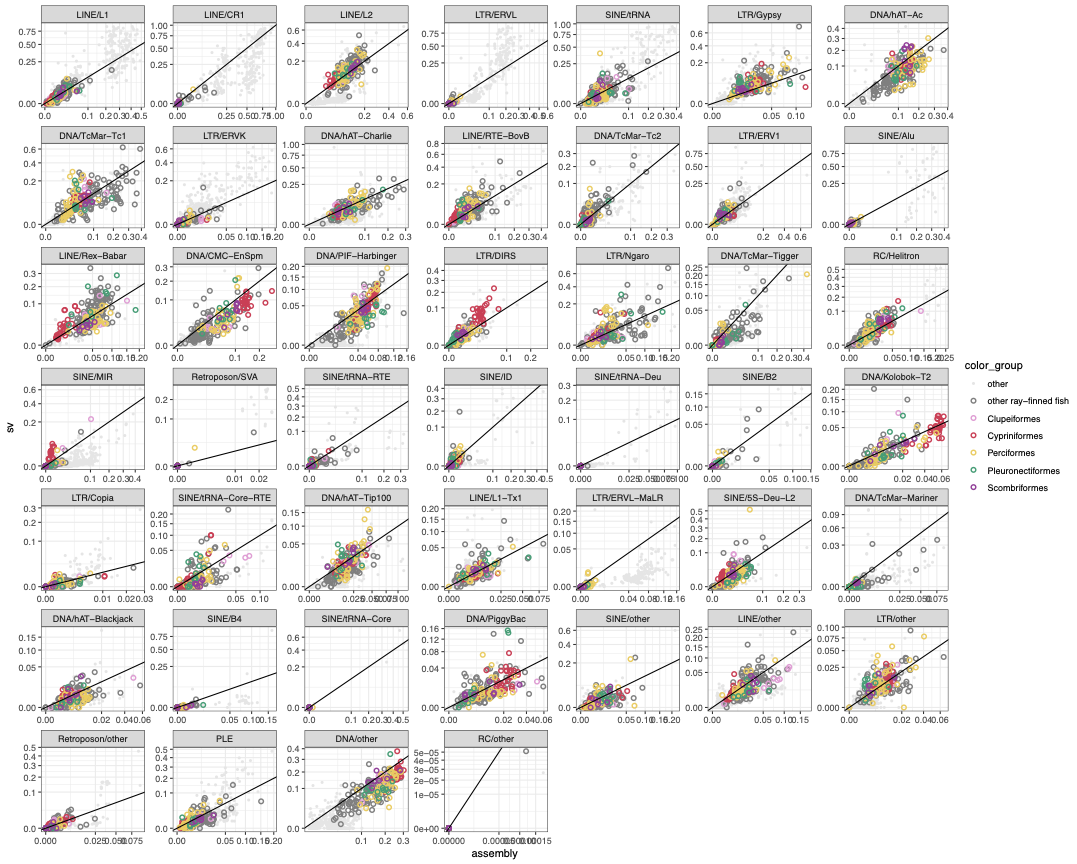

**Figure S4 -** The abundance of all major TE families in SVs (y-axis) and in genome assemblies (x-axis) across ray-finned fishes. TE abundance is quantified by their proportion among all classified TEs. Assemblies are colored by major ray-finned fish orders. A one-to-one line is drawn to indicate the expected distribution if TE composition is identical between SVs and assemblies.

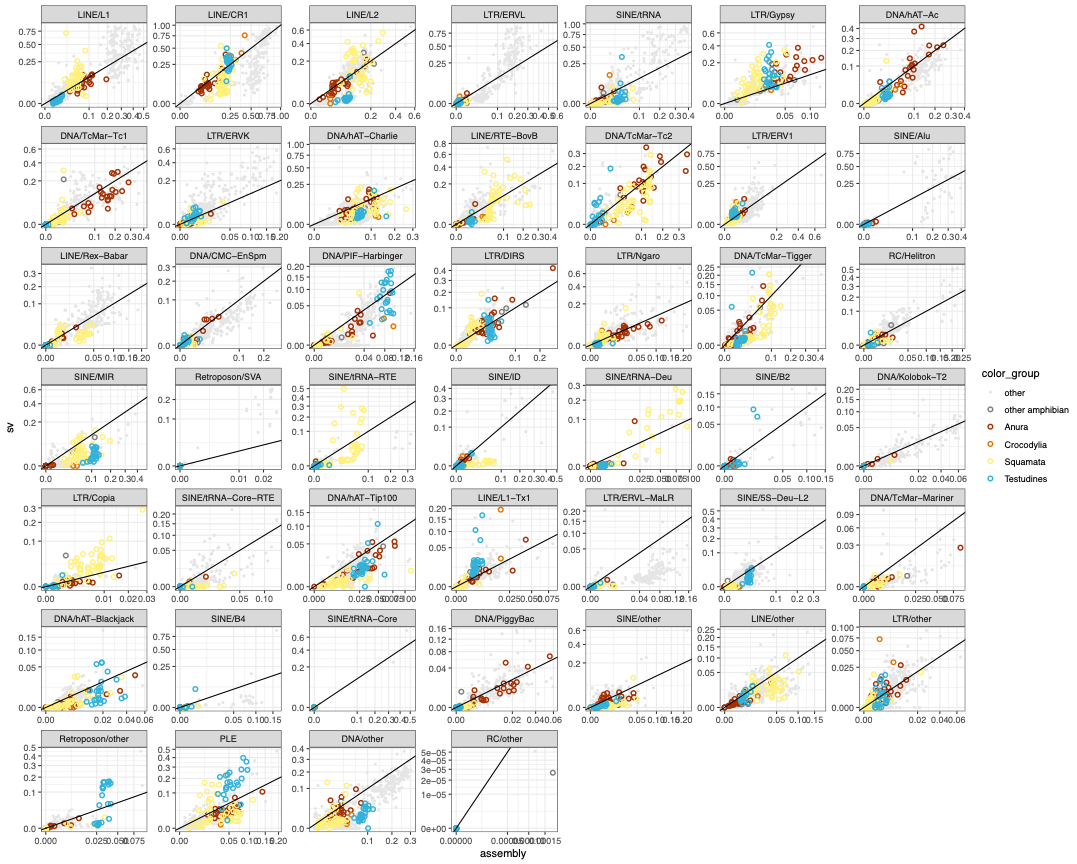

**Figure S5 -** The abundance of all major TE families in SVs (y-axis) and in genome assemblies (x-axis) across amphibians and reptiles. TE abundance is quantified by their proportion among all classified TEs. Assemblies are colored by major amphibian and reptile orders. A one-to-one line is drawn to indicate the expected distribution if TE composition is identical between SVs and assemblies.

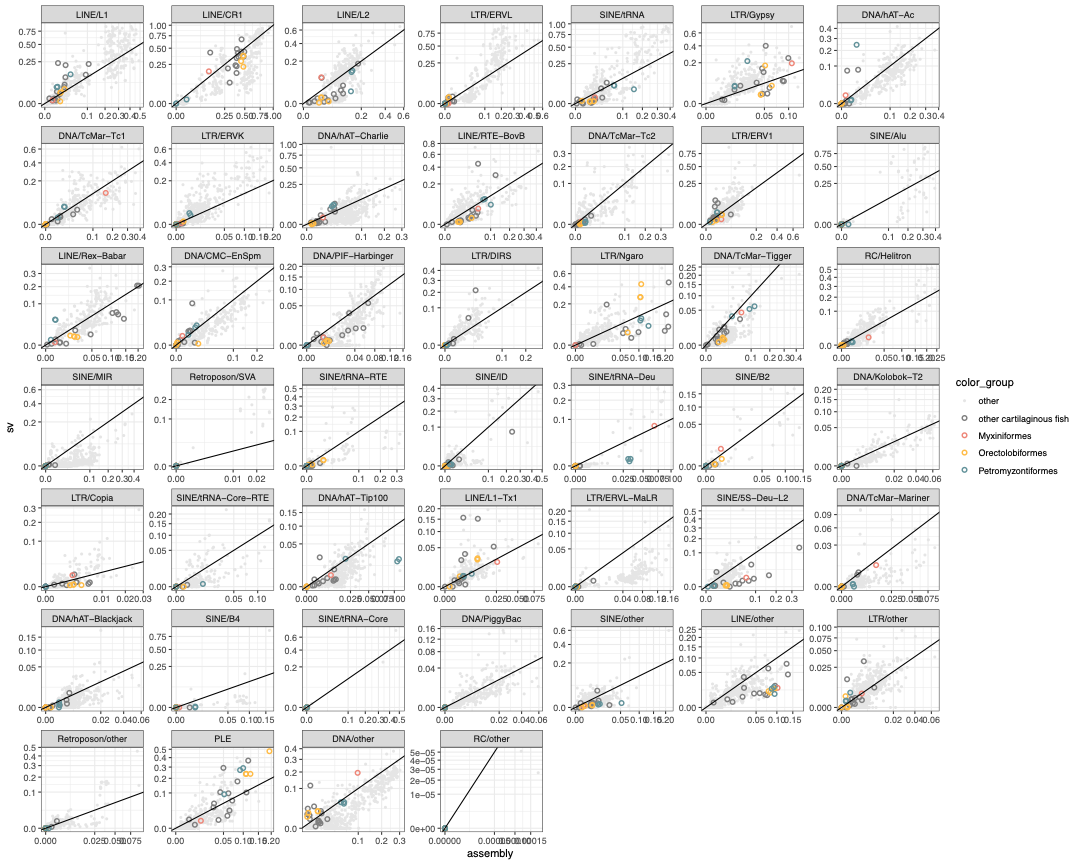

**Figure S6 -** The abundance of all major TE families in SVs (y-axis) and in genome assemblies (x-axis) across cartilaginous fishes. TE abundance is quantified by their proportion among all classified TEs. Assemblies are colored by major cartilaginous fish orders. A one-to-one line is drawn to indicate the expected distribution if TE composition is identical between SVs and assemblies.
